# Genetic analysis of pyrimidine biosynthetic enzymes in *Plasmodium falciparum*

**DOI:** 10.1101/2025.11.30.691290

**Authors:** Krithika Rajaram, Montana L. Sievert, Rubayet Elahi, Lucas B. Dillard, James Blauwkamp, Sabrina Absalon, Sean T. Prigge

**Affiliations:** Department of Molecular Microbiology and Immunology, Johns Hopkins Bloomberg School of Public Health, Baltimore, MD; Department of Microbiology, The Ohio State University, Columbus, OH; Department of Biophysics and Biophysical Chemistry, Johns Hopkins University School of Medicine, Baltimore, MD; Department of Biochemistry, Molecular Biology and Toxicology, Indiana University School of Medicine, Indianapolis, IN

## Abstract

The malaria parasite *Plasmodium falciparum* depends entirely on *de novo* pyrimidine synthesis, as it is unable to salvage these essential nucleotides. This reliance makes the pyrimidine biosynthesis pathway a compelling target for antimalarial drugs, with several inhibitors targeting its rate-limiting enzyme, dihydroorotate dehydrogenase (*Pf*DHODH), already in clinical development. In this study, we investigated the roles of three other pathway enzymes – aspartate transcarbamoylase (*Pf*ATC), carbamoyl phosphate synthetase II (*Pf*CPSII), and dihydroorotase (*Pf*DHO). *Pf*ATC features a unique N-terminal extension predicted to serve as an apicoplast trafficking peptide. However, using antibodies against the native protein and an epitope-tagged version, we found no evidence of apicoplast localization. Knockdown of *Pf*ATC expression proved lethal and could not be rescued by an apicoplast metabolic bypass. Complementation assays further revealed that truncation of the N-terminal domain impaired parasite growth, suggesting that this region is important for *Pf*ATC function or stability *in vivo*. *Pf*CPSII, which harbors large *Plasmodium*-specific insertions between its catalytic domains, was likewise found to be essential for parasite proliferation. To assess the role of *Pf*DHO, we engineered parasites to salvage uracil via heterologous expression of a yeast enzyme. Deletion of *Pf*DHO in this parasite line resulted in uracil auxotrophy, confirming the enzyme’s essential function in pyrimidine synthesis. Together, these findings reveal multiple vulnerable nodes within the pyrimidine biosynthesis pathway.

**AUTHOR SUMMARY:** Nucleotides are central metabolites that serve as building blocks for DNA and RNA, act as key energy carriers, and function as cofactors or regulators in several metabolic pathways. To satisfy these diverse demands, most organisms rely on both nucleotide salvage and *de novo* synthesis. The malaria parasite *Plasmodium falciparum* acquires purine nucleotides from the host but lacks the capacity to salvage pyrimidines, making *de novo* pyrimidine synthesis essential. Several enzymes in this pathway differ from their human counterparts in sequence, domain architecture, and evolutionary origin, enhancing their potential as selective drug targets. Dihydroorotate dehydrogenase (PfDHODH), the fourth enzyme in the pathway, has already been validated as an antimalarial target. Here, we systematically examined upstream enzymes using molecular genetic approaches. Each proved essential for asexual blood-stage parasite survival, with the *Plasmodium*-specific N-terminal extension of aspartate carbamoyltransferase (*Pf*ATC) required for optimal growth. The introduction of a yeast uracil salvage enzyme rescued parasites depleted of these biosynthetic enzymes, demonstrating that their essential functions are confined to pyrimidine production and that their distinctive structural features do not support additional metabolic roles. In summary, these results delineate additional enzymatic steps in this important metabolic pathway that warrant continued investigation from both biological and translational angles.

## INTRODUCTION

The transition from a free-living to an obligate parasitic lifestyle is marked by a reduction of core metabolic pathways and the acquisition of nutrient salvage mechanisms, leading to irreversible dependence on the host. Apicomplexan parasites, which cause a range of human and animal diseases, have undergone genomic streamlining to adapt to their host niches, enabling them to salvage several key building blocks like amino acids, lipids, and nucleotides (1–3). While all apicomplexans rely on purine salvage, their capacity for pyrimidine uptake varies (4–6). *Plasmodium falciparum*, the causative agent of human malaria, is unable to use exogenous pyrimidines and requires *de novo* synthesis, making this pathway an attractive target for therapeutic intervention (7–11).

Clinical symptoms of malaria stem from the rapid proliferation of *P. falciparum* within host red blood cells, which, being anucleate, have low nucleotide demands and limited pyrimidine availability (12). The parasite sustains its high metabolic needs by synthesizing the pyrimidine uridine 5’-monophosphate (UMP) through a conserved six-step pathway that begins with carbamoyl phosphate synthetase II (CPSII) (**Fig. 1**) (13, 14). CPSII catalyzes the formation of carbamoyl phosphate from bicarbonate and glutamine (15, 16). Carbamoyl phosphate then undergoes condensation with aspartate in a reaction catalyzed by aspartate transcarbamoylase (ATC) to yield carbamoyl aspartate and inorganic phosphate (17–19).

**Fig. 1.**
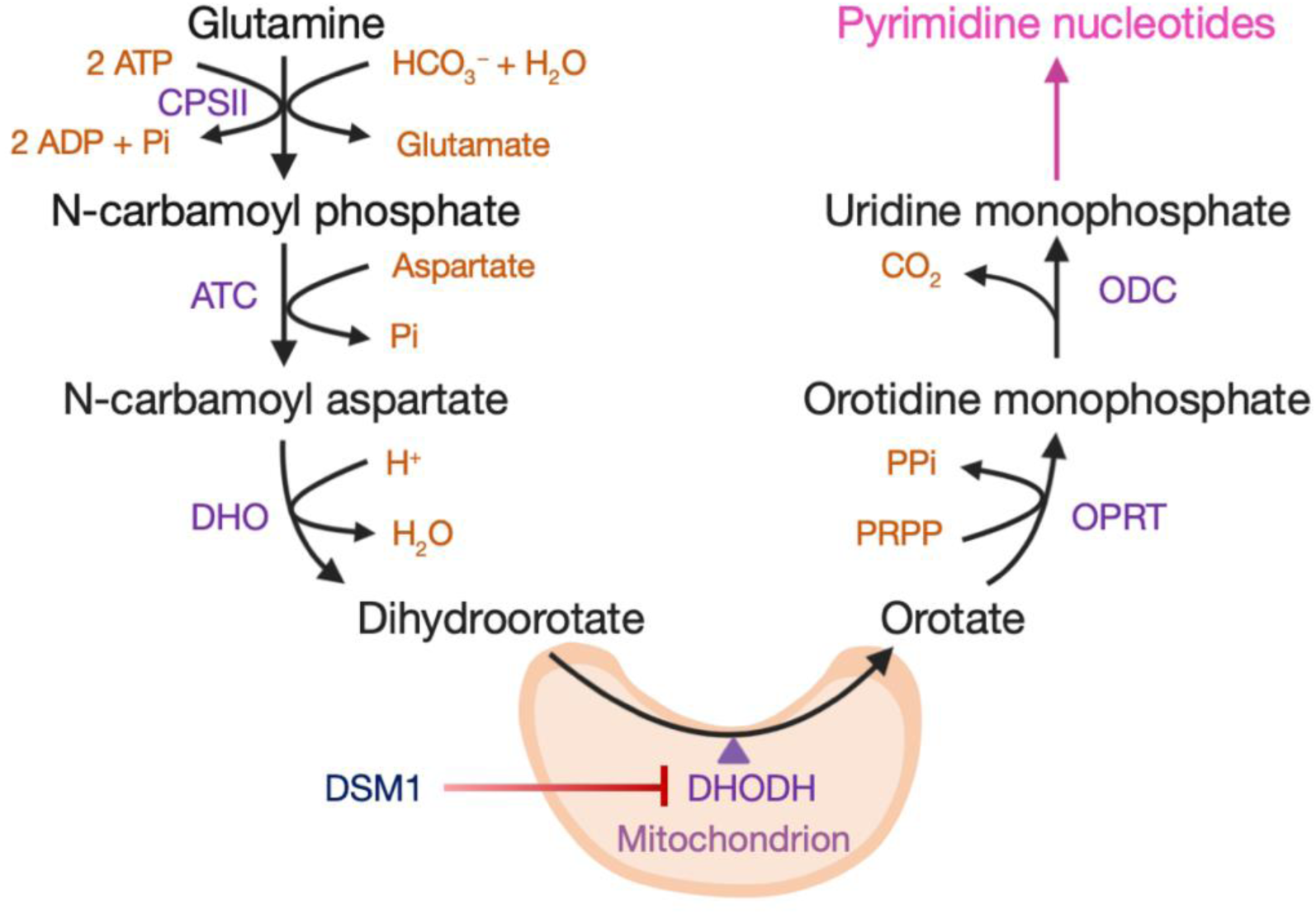
Pyrimidine biosynthesis in *P. falciparum*. This figure illustrates the enzymatic reactions involved in the *de novo* biosynthesis of pyrimidines in *P. falciparum*. The pathway begins with the conversion of glutamine and bicarbonate into carbamoyl phosphate by the enzyme carbamoyl phosphate synthetase II (*Pf*CPSII). Subsequent steps catalyzed by aspartate transcarbamoylase (*Pf*ATC) and dihydroorotase (*Pf*DHO) lead to the formation of dihydroorotate. Dihydroorotate is then oxidized to orotate by dihydroorotate dehydrogenase (*Pf*DHODH), a mitochondrial enzyme that can be inhibited by a class of triazolopyrimidine compounds including DSM1 (20). Orotate is subsequently converted to orotidine-5’-monophosphate (OMP) by orotate phosphoribosyltransferase (*Pf*OPRT), and OMP is decarboxylated to uridine 5’-monophosphate (UMP) by orotidine-5’-monophosphate decarboxylase (*Pf*ODC). UMP is a precursor for the synthesis of all pyrimidine nucleotides, which are essential for DNA and RNA synthesis.

Dihydroorotase (DHO) catalyzes the next reaction in the pathway, converting carbamoyl aspartate to dihydroorotate (21, 22). Dihydroorotate is oxidized to orotate by dihydroorotate dehydrogenase (DHODH), a flavin-dependent mitochondrial enzyme that uses ubiquinone as a terminal electron acceptor in malaria parasites (23). *P. falciparum* DHODH (*Pf*DHODH) activity is thought to be the primary driver for maintaining mitochondrial electron transport chain (mETC) function in asexual blood-stage parasites, which primarily rely on glycolysis for ATP production (24–26). The role of the mETC during this stage appears to be largely limited to the regeneration of ubiquinone for *Pf*DHODH (24). Consequently, mETC inhibitors like atovaquone are ineffective against *P. falciparum* lines engineered to express a cytosolic yeast DHODH homolog that uses fumarate instead of ubiquinone (24, 27–30).

Orotate phosphoribosyltransferase (OPRT) performs the penultimate step in the pathway, transferring a ribose 5-phosphate group from phosphoribosyl pyrophosphate to orotate to form orotidine 5′-monophosphate (OMP) (23, 31, 32). OMP is then converted to the pathway product UMP by OMP decarboxylase (ODC) (23, 32). UMP is subsequently converted into other pyrimidine and deoxypyrimidine nucleotides through a series of enzymatic steps (7, 13).

Phylogenetic analyses reveal that the pyrimidine biosynthetic enzymes in *P. falciparum* have mixed evolutionary origins: *Pf*CPSII, *Pf*DHO, and *Pf*OPRT are more closely related to prokaryotic orthologs, while the remaining enzymes exhibit features of both bacterial and eukaryotic lineages (11, 13). *Pf*CPSII and *Pf*ATC are also unusual in that they contain large insertions of unknown function (14, 19, 33). Beyond these sequence-level distinctions, the structural organization of the pathway also differs markedly from that of the human host (11, 13). In humans and other multicellular eukaryotes, the first three steps of pyrimidine biosynthesis are catalyzed by a single multifunctional protein, CAD, which combines the activities of CPSII, ATC, and DHO (34). Similarly, OPRT and ODC are fused into a bifunctional enzyme known as UMP synthase in many eukaryotes (32, 35). In *P. falciparum*, however, *Pf*OPRT and *Pf*ODC are expressed as separate proteins that form a heterotetrameric complex (23, 36–40). The distinct architecture and evolutionary divergence of the *Plasmodium* enzymes from their human counterparts further enhance their potential as selective drug targets. Consistent with this, gene essentiality screens in *P. falciparum* and *P. berghei* indicate that most of these enzymes are required for parasite survival or associated with significant fitness costs (**Table 1**) (41). Notably, *Pf*DHODH, the only clinically validated target in this pathway, has already been the focus of extensive antimalarial drug development efforts (20, 42–50).

**Table 1.** Essentiality of pyrimidine biosynthesis pathway genes from genetic screens in *Plasmodium*. a – Zhang et al., 2018 (41); b – Schwach et al., 2015 (51); c – Oberstaller et al., 2025 (52); d – Elsworth et al., 2025 (53)

| Gene ID (PF3D7_) | Gene name | Essentiality in asexual blood-stage *Plasmodium* |  |  |  |  |
| --- | --- | --- | --- | --- | --- | --- |
|  |  | *P. falciparum* (a) |  | *P. berghei* (b)** | *P. knowlesi* (c, d) |  |
|  |  | MFS* | MIS* |  | MIS+<sup>#</sup> | HMS<sup>##</sup> |
| 1308200 | Carbamoyl phosphate synthetase II (CPSII) | -2.989 | 0.119 | Essential | 0.0035 | 0.06 |
| 1344800 | Aspartate transcarbamoylase (ATC) | -2.437 | 0.408 | Essential | 0.0017 | 0.28 |
| 1472900 | Dihydroorotase (DHO) | -1.259 | 1.0 | No data available | 0.0035 | 0.18 |
| 0603300 | Dihydroorotate dehydrogenase (DHODH) | -0.3099 | 0.125 | Slow | 0.012 | 0.18 |
| 0512700 | Orotate phosphoribosyltransferase (OPRT) | -3.229 | 0.214 | Slow | 0.0257 | 0.13 |
| 1023200 | Orotidine 5'-monophosphate decarboxylase (OMPDC) | -2.664 | 0.614 | Slow | 0.001 | 0.18 |

\* The genome-wide screen for essential genes in *P. falciparum* employed the mutagenesis fitness score (MFS; ranging from -4.5 to 2) and mutagenesis index score (MIS; ranging from 0 to 1), to assess gene mutability (41). Low MFS and MIS scores are indicative of gene essentiality.

In this study, we systematically dissected the pyrimidine biosynthetic pathway in *P. falciparum*, using gene knockdown and metabolic bypass approaches to investigate the roles of multiple enzymes. Although *Pf*CPSII and *Pf*ATC possess unique sequence features absent from host counterparts, and DHODH has been reported to perform an additional function in *Toxoplasma gondii* (54), our findings demonstrate that these enzymes, along with *Pf*DHO, play a singular, essential role in pyrimidine synthesis during the asexual blood stage. By defining the essential metabolic function of these parasite enzymes, our work expands the pool of candidate antimalarial targets within this important biosynthetic pathway.

## RESULTS

### Subcellular location of *P. falciparum* aspartate transcarbamoylase (*Pf*ATC)

The ATC gene in *P. falciparum* encodes a 375-amino acid (aa) protein (*Pf*ATC) that forms a homotrimer. Unlike its *E. coli* counterpart, the functional *Pf*ATC complex lacks regulatory subunits, suggesting it is not subject to feedback inhibition by the pathway’s end-product, UMP (18, 19, 33). *Pf*ATC features an N-terminal extension that is conserved among *Plasmodium* species but absent in orthologs from animals, fungi, and bacteria (**Fig. 2A**). This domain is predicted to function as an apicoplast trafficking peptide based on analyses from three different bioinformatic tools: PATS, ApicoAP, and PlasmoAP (55–57). A previous attempt to determine the subcellular location of *Pf*ATC showed that a GFP-tagged copy of the protein localized to a region distinct from the endoplasmic reticulum (ER); however, since the study did not include an apicoplast-specific marker, the precise location of *Pf*ATC remains unresolved (19).

**Fig. 2.**
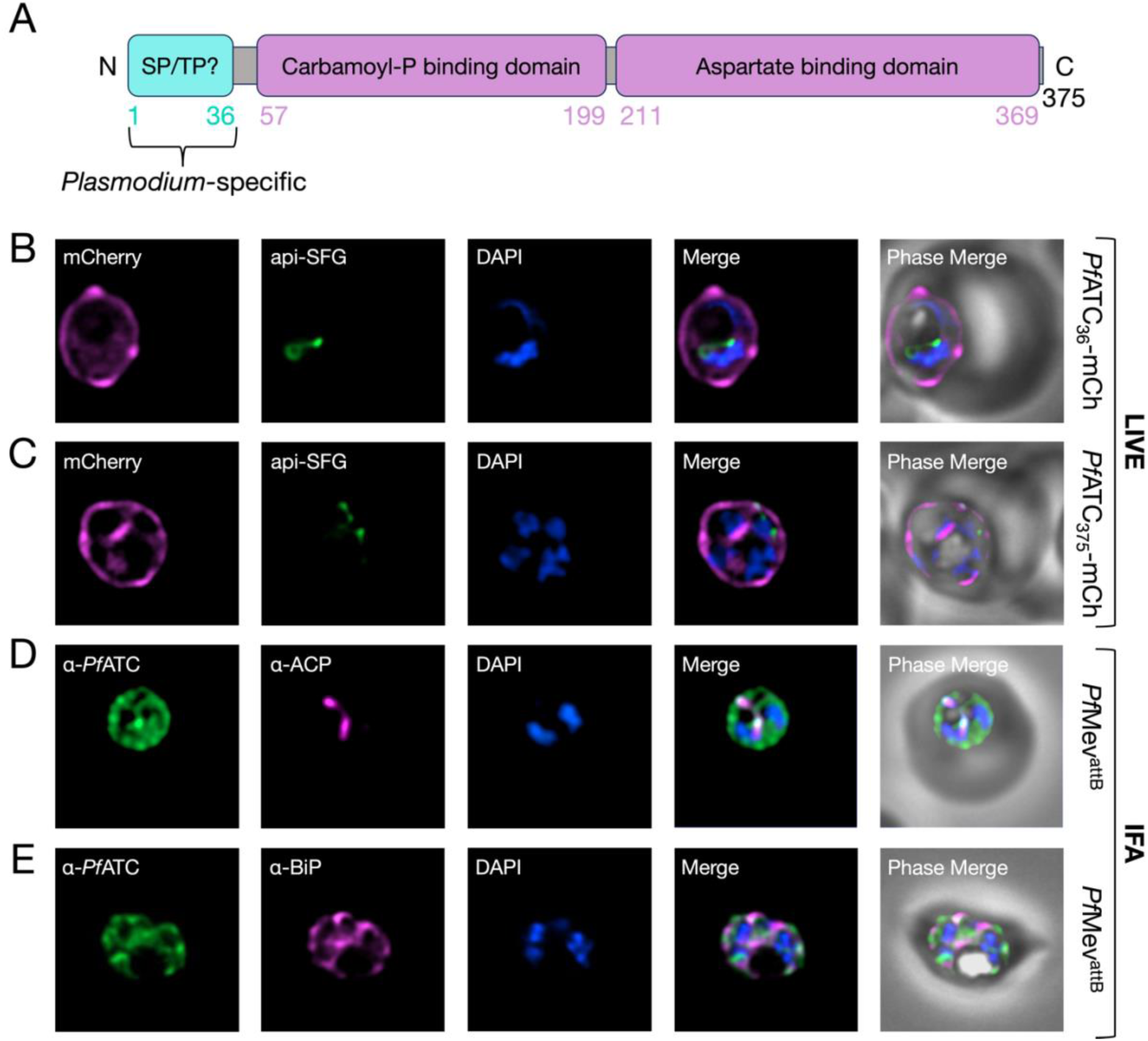
*Pf*ATC does not localize to the apicoplast. (A) Predicted (cyan) and annotated (purple) functional domains of the *Pf*ATC protein are depicted. The first 36 amino acids of *Pf*ATC are predicted to form a signal peptide (SP) and an apicoplast transit peptide (TP). (B-C) Representative live fluorescence images of parasites expressing (B) truncated or (C) full-length *Pf*ATC fused to mCherry (mCh) showed that the tagged proteins are mostly peripherally located and do not colocalize with the apicoplast marker api-SFG. (D-E) Representative images from immunofluorescence assays (IFA) on parental *Pf*Mev^attB^ parasites probed with α-*Pf*ATC and (D) apicoplast (α-ACP), or (E) ER-specific (α-BiP) antibodies did not reveal substantial overlap of signals. Quantification by Manders’ coefficient M1 (green signal within red signal) yielded a value of 0.138 (±0.098; n = 19) for α-*Pf*ATC and α-ACP, and 0.372 (±0.221; n = 19) for α-*Pf*ATC and α-BiP. DAPI (blue) stains the parasite nucleus. Images represent fields that are 10 μm long by 10 μm wide.

To determine if *Pf*ATC is indeed an apicoplast protein, we fused a C-terminal mCherry tag to the first 36-aa of *Pf*ATC (*Pf*ATC_36_-mCh) which corresponds to the predicted apicoplast trafficking peptide. The fusion protein was expressed from a plasmid integrated into an *att*B site in *Pf*Mev^attB^ parasites (**Fig. S1A-C**) (58, 59). This parasite line features a built-in fluorescent apicoplast reporter consisting of the apicoplast trafficking peptide of acyl carrier protein fused to superfolder GFP (api-SFG). Live microscopy revealed that *Pf*ATC_36_-mCh localizes primarily to the parasite periphery, along with a weaker cytoplasmic signal that does not overlap with the apicoplast (**Fig. 2B**). Expression of full-length ATC fused to mCherry (*Pf*ATC_375_-mCh) also displayed a similar localization pattern, showing no overlap with api-SFG (**Fig. 2C, Fig. S1D**).

To assess the location of endogenous *Pf*ATC, we generated antibodies against recombinant *Pf*ATC protein (**Fig. S2**). Immunofluorescence imaging of parasites using the α-*Pf*ATC antibodies revealed predominant staining in the parasite cytoplasm, with some peripheral signal (**Fig. 2D-E**). Co-labeling with antibodies against apicoplast or ER markers showed minimal overlap in signals (**Fig. 2D-E**). Attempts to obtain higher-resolution imaging data using expansion microscopy were unsuccessful since the α-*Pf*ATC antibodies produced non-specific staining in expanded cells. Overall, our findings indicate that *Pf*ATC is not located in the apicoplast, but it may be present in more than one subcellular location.

### *Pf*ATC is essential in blood-stage malaria parasites

Given its role in pyrimidine synthesis, we anticipated that blocking *Pf*ATC expression would be lethal to blood-stage malaria parasites. Consistent with this, *Pf*ATC was assigned a high essentiality score in a forward genetic screen in *P. falciparum* (**Table 1**) (41). For further validation, we employed a TetR-aptamer system to conditionally knock down *Pf*ATC expression (60, 61). A linearized pKD plasmid carrying TetR-DOZI components was introduced at the 3’ end of the *Pf*ATC gene in *Pf*Mev^attB^ parasites using CRISPR/Cas9-mediated gene editing (**Fig. S3A-B**) (62). The modified *Pf*ATC gene encodes a C-terminal 2×FLAG tag for protein visualization, and a 10x aptamer array in its 3’ UTR so the transcribed mRNA can be regulated by the TetR-DOZI module and its ligand anhydrotetracycline (aTet). The genotype of the conditional knockdown (*Pf*ATC^CD^) parasite line was confirmed by PCR (**Fig. S3C**). The parasites were maintained continuously in medium containing aTet to allow translation of *Pf*ATC.

To determine if *Pf*ATC is an essential protein, parasites were washed free of aTet to knock down *Pf*ATC expression, and their growth was monitored for 8 days. Immunoblotting with an α-FLAG antibody revealed strong expression of FLAG-tagged *Pf*ATC in parasite lysates on Day 0, which gradually decreased to undetectable levels by Day 3 (**Fig. 3A**). By Day 4, parasites cultured without aTet displayed a significant growth defect (**Fig. 3B**).

**Fig. 3.**
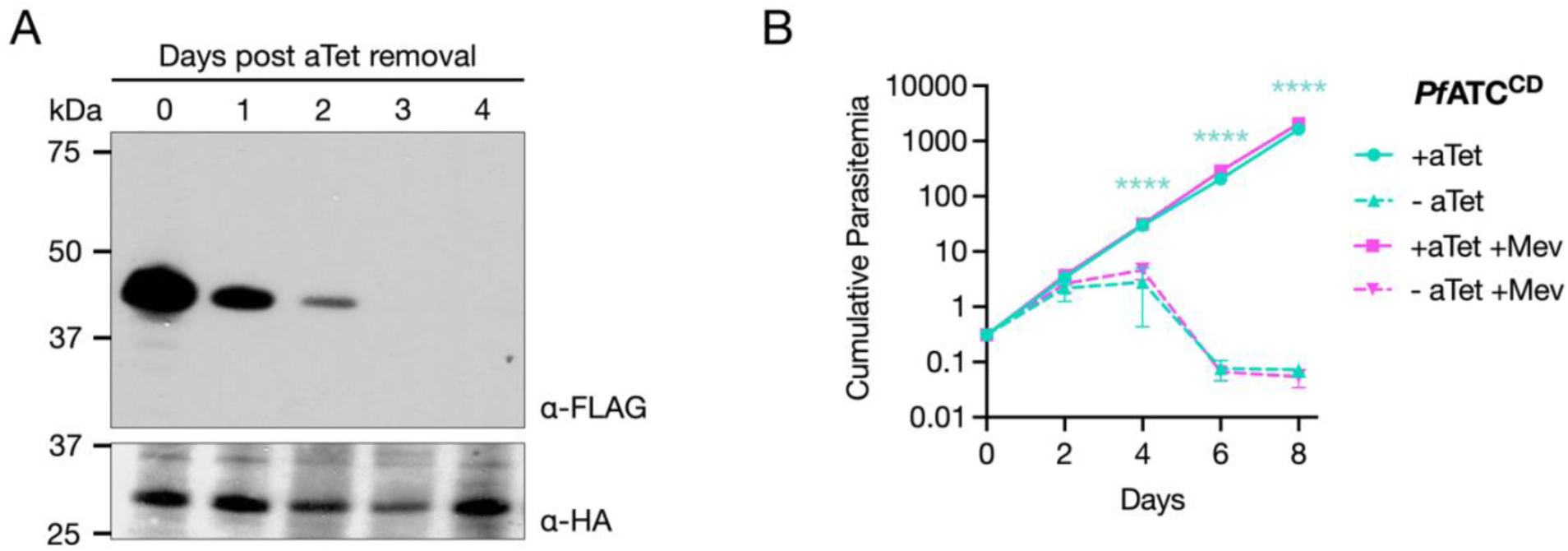
*Pf*ATC is essential for the survival of blood-stage *P. falciparum* parasites. (A) Western blot analysis using an α-FLAG antibody to probe for *Pf*ATC (expected MW: 45 kDa) revealed efficient knockdown of expression following aTet removal from *Pf*ATC^CD^ parasite cultures. An α-hemagglutinin (HA) antibody was used to detect HA-tagged api-SFG (expected MW of processed form: 28.1 kDa), which served as a loading control. (B) Growth of *Pf*ATC^CD^ parasites in the presence or the absence of aTet and mevalonate (Mev) was monitored by flow cytometry for 8 days. Removal of aTet led to parasite death, which could not be rescued by Mev addition. Two biological experiments were conducted in quadruplicate. Cumulative parasitemia values were plotted on a logarithmic Y-axis. Error bars represent standard deviations from the means. Statistical analysis was conducted using GraphPad Prism 10 (GraphPad Software, Inc). Two-way ANOVA was followed by Bonferroni’s correction (****, P ≤ 0.0001; significant differences are shown for pairwise comparisons of the - aTet and - aTet +Mev conditions relative to the +aTet control).

Parasites with apicoplast defects in the *Pf*Mev^attB^ background can be rescued by the addition of mevalonate, which is converted into essential isoprenoid building blocks through an engineered cytosolic pathway (58). To determine if an apicoplast bypass could partially or fully rescue *Pf*ATC-depleted parasites, a fraction of the aTet-free *Pf*ATC^CD^ culture was split into media containing mevalonate on Day 0 of the growth assay. The addition of mevalonate did not alter the kinetics of parasite death (**Fig. 3B**). From these data, we conclude that *Pf*ATC is an essential protein whose function cannot be complemented by a metabolic bypass for the apicoplast. This finding, together with the observed localization pattern for *Pf*ATC (**Fig. 2**), suggests that the enzyme plays an essential role outside of the apicoplast.

### N-terminal truncation of *Pf*ATC impacts parasite growth

Since the N-terminal extension of *Pf*ATC is an uncommon feature for aspartate transcarbamoylases (**Fig. 2A**), we asked if its removal would have any impact on the protein’s function in *P. falciparum* parasites. An mCherry tag was fused to the C-terminus of truncated *Pf*ATC missing the first 36-aa (*Pf*ATC_Δ36_-mCh). The fusion protein was expressed from a chromosomally integrated plasmid in the *Pf*ATC^CD^ parasite line (**Fig. S4A**). A control parasite line was generated by introducing a second copy of full-length *Pf*ATC (*Pf*ATC_375_-mCh) into *Pf*ATC^CD^ parasites (**Fig. S4B**). Live microscopy of *Pf*ATC_Δ36_-mCh parasites revealed diffuse mCherry signal throughout the parasite cytoplasm as opposed to the peripheral signal observed with *Pf*ATC_375_-mCh (**Fig. 4A-B**). These results demonstrate that the N-terminus of *Pf*ATC affects protein localization.

**Fig. 4.**
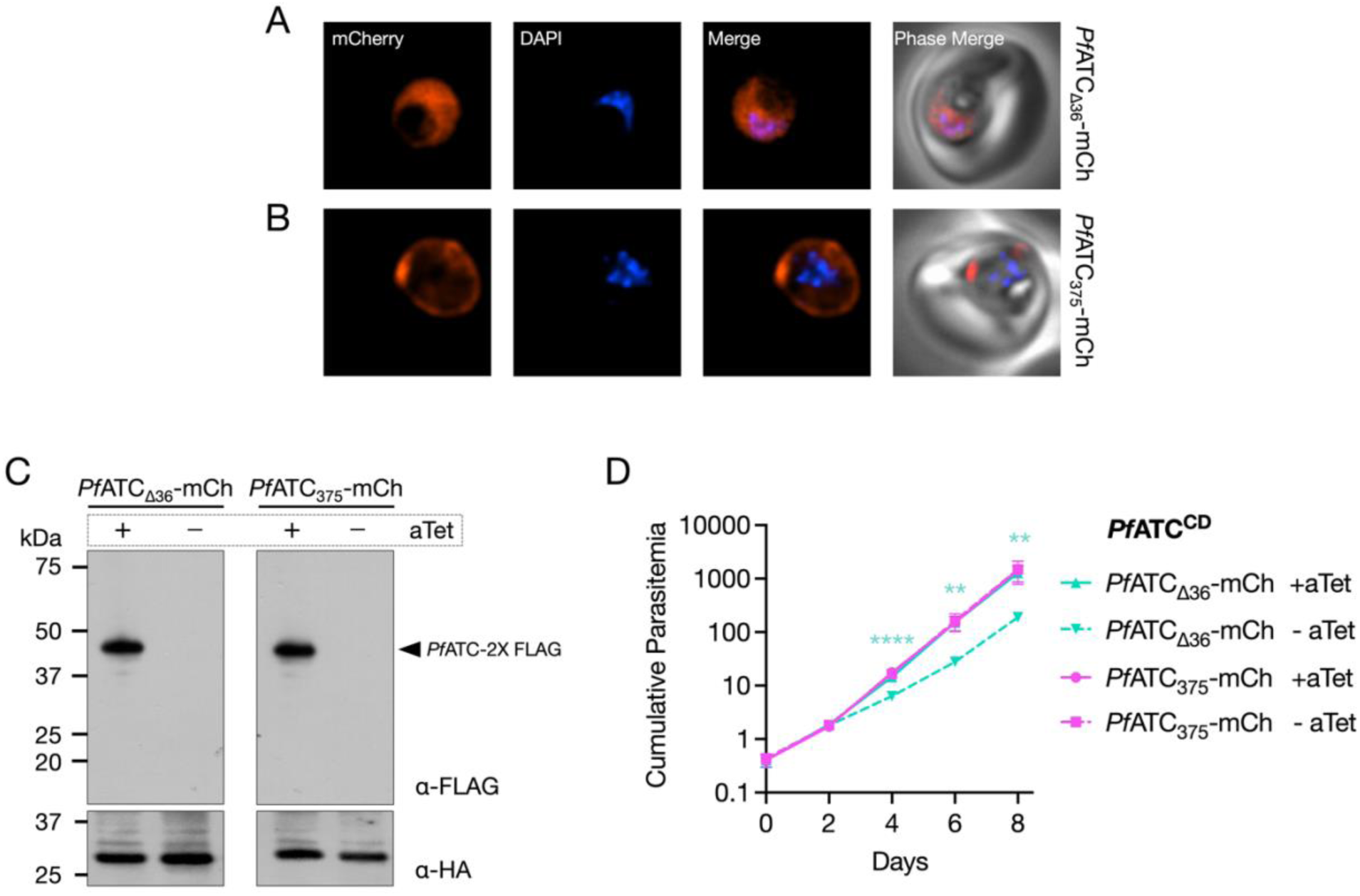
N-terminal truncation of *Pf*ATC impacts its localization and parasite growth. Representative live fluorescence microscopy images of *Pf*ATC^CD^ parasites expressing (A) truncated (*Pf*ATC_Δ36_-mCh) or (B) full-length (*Pf*ATC_375_-mCh) constructs of *Pf*ATC fused to mCherry showed that N-terminal truncation resulted in a shift from peripheral localization to diffuse cytoplasmic distribution of *Pf*ATC. Images represent fields that are 10 μm long by 10 μm wide. (C) Western blot analysis using an α-FLAG antibody on parasite lysates collected from Day 0 (+) and Day 8 (-) of aTet washout revealed efficient knockdown of endogenous FLAG-tagged *Pf*ATC (45 kDa). The α-HA antibody detecting HA-tagged api-SFG (expected MW of processed form: 28.1 kDa) was used as a loading control. (D) Growth of *Pf*ATC^CD^ parasites in the absence of aTet was entirely rescued by second-copy expression of *Pf*ATC_375_-mCh, and only partially rescued by *Pf*ATC_Δ36_-mCh. Two independent biological experiments were performed in quadruplicate. Cumulative parasitemia values were plotted on a logarithmic Y-axis. Error bars represent standard deviations from the mean. Statistical analysis was conducted using GraphPad Prism 10 (GraphPad Software, Inc). Two-way ANOVA was followed by Bonferroni’s correction (**, P ≤ 0.01; ****, P ≤ 0.0001; asterisks indicate significant differences between +aTet and - aTet *Pf*ATC_Δ36_-mCh conditions).

Next, we examined if truncated *Pf*ATC could rescue parasites depleted of endogenous *Pf*ATC. The *Pf*ATC^CD^ parasites expressing truncated or full-length *Pf*ATC-mCh were cultured in media with or without aTet, and their growth was monitored over a period of 8 days. Successful knockdown of endogenous *Pf*ATC was confirmed in cultures lacking aTet (**Fig. 4C**). Parasites expressing *Pf*ATC_375_-mCh grew normally even without aTet, demonstrating that the full-length fusion protein can fully complement native *Pf*ATC function (**Fig. 4D**). By contrast, expression of *Pf*ATC_Δ36_-mCh only partially rescued growth, with nearly a 10-fold reduction in cumulative parasitemia on days 6 and 8 of the growth assay (**Fig. 4D**). Taken together, our results show that removing the N-terminal of *Pf*ATC impacts its subcellular location and parasite fitness.

### *Pf*CPSII and *Pf*DHO are required for parasite survival

The substrate for ATC, carbamoyl phosphate, is synthesized by carbamoyl phosphate synthetase II (CPSII). ATC then converts it into carbamoyl aspartate, which is subsequently utilized by dihydroorotase (DHO) (**Fig. 1**). The *P. falciparum* CPSII (*Pf*CPSII) gene encodes a 2375-aa product with long insertions between putative functional domains, making it one of the largest known CPS family enzymes (**Fig. 5A**) (14). *Pf*CPSII was predicted to be essential in the forward genetic screen, whereas *Pf*DHO was unexpectedly deemed dispensable (**Table 1**) (41). Since neither gene has been characterized in depth, we generated conditional knockdown lines for *Pf*CPSII (*Pf*CPSII^CD^) and *Pf*DHO (*Pf*DHO^CD^) in the *Pf*Mev^attB^ background to determine their essentiality in blood-stage malaria parasites (**Fig. S5A-C**). As before, the endogenous genes were modified to encode C-terminal FLAG tags for tracking protein fate. Immunofluorescence microscopy indicated that *Pf*CPSII-FLAG localizes to the cytoplasm (**Fig. 5B**). Unfortunately, *Pf*DHO could not be reliably detected.

**Fig. 5.**
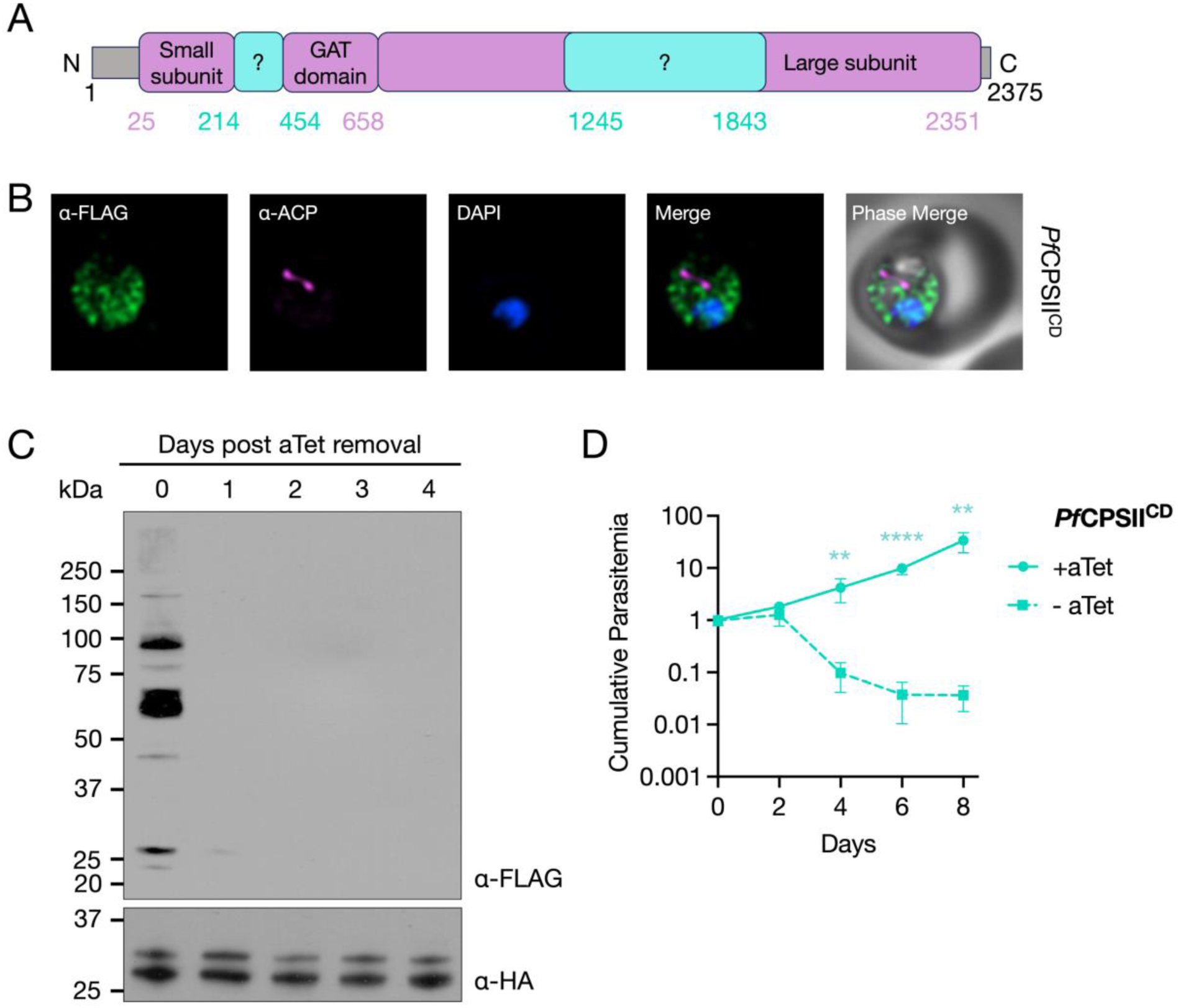
*Pf*CPSII is an essential protein in blood-stage *P. falciparum* parasites. (A) Insertions (cyan) and annotated (purple) functional domains of the *Pf*CPSII protein are depicted. GAT – glutamine amidotransferase domain. (B) Representative images from immunofluorescence assays (IFA) on *Pf*CPSII^CD^ parasites probed with α-FLAG and α-ACP (apicoplast) antibodies indicated that endogenous FLAG-tagged CPSII localized to the parasite cytoplasm. Images represent fields that are 10 μm long by 10 μm wide. (C) Western blot analysis using an α-FLAG antibody to probe for *Pf*CPSII revealed efficient knockdown of expression following aTet removal from *Pf*CPSII^CD^ parasite cultures. The α-HA antibody detecting HA-tagged api-SFG was used as a loading control. The two bands indicate precursor (33 kDa) and mature (28.1 kDa) forms of the apicoplast protein. (D) Growth of *Pf*CPSII^CD^ parasites in the presence or the absence of aTet was measured by flow cytometry for 8 days. Removal of aTet led to parasite death. Two biological experiments were conducted in quadruplicate. Cumulative parasitemia values were plotted on a logarithmic Y-axis. Error bars represent standard deviations from the mean. Statistical analysis was conducted using GraphPad Prism 10 (GraphPad Software, Inc). Two-way ANOVA was followed by Bonferroni’s correction (**, P ≤ 0.01; ****, P ≤ 0.0001).

The transgenic parasites were divided into media with and without aTet and monitored for 8 days. The levels of *Pf*CPSII in the *Pf*CPSII^CD^ parasites were followed by immunoblotting with an α-FLAG antibody. While the expected size of *Pf*CPSII is ∼273 kDa, we did not detect a protein at this molecular weight. Instead, we observed several smaller bands on Day 0 that were no longer detectable in parasites after aTet removal, suggesting that they are processed forms of *Pf*CPSII (**Fig. 5C**). The aTet-depleted *Pf*CPSII^CD^ parasites exhibited a pronounced growth defect by Day 4, confirming that *Pf*CPSII is required for blood-stage development of *P. falciparum* (**Fig. 5D**).

The *Pf*DHO^CD^ parasites did not show appreciable reduction in *Pf*DHO protein levels upon aTet removal and displayed only modest fitness defects under these conditions (**Fig. S5D-E**). Since the knockdown was not very effective, we attempted to knock out the *Pf*DHO gene in *Pf*Mev^attB^ parasites using CRISPR/Cas9. Despite multiple attempts, we failed to generate a deletion line, suggesting that *Pf*DHO may be essential.

To further investigate gene essentiality, we repeated the *Pf*DHO deletion experiments in the presence of a controllable pyrimidine bypass system. We introduced the *Saccharomyces cerevisiae* FUR1 gene (uracil phosphoribosyltransferase), tagged at the N-terminus with mCherry (mCh-FUR1), into the *att*B locus of *Pf*Mev^attB^ parasites to create the *Pf*Mev^FUR1^ line (**Fig. S6A**) (63, 64). FUR1 is expected to convert supplemented uracil into UMP, the end-product of *de novo* pyrimidine biosynthesis (**Fig. 6A**). Live microscopy confirmed the expression of mCh-FUR1 in the parasite cytoplasm (**Fig. 6B**). To test the functionality of FUR1, we asked whether uracil supplementation could rescue *Pf*Mev^FUR1^ parasites from growth inhibition caused by DSM1, a drug that targets the fourth enzyme of the *de novo* pathway, *Pf*DHODH (**Fig. 1**, **Fig. 6C**). Control parasites not exposed to DSM1 grew equally well with or without 50 μM uracil, indicating that uracil addition alone does not affect parasite growth. As expected, DSM1 treatment inhibited parasite growth, which was fully rescued by uracil, confirming that the FUR1 bypass is functional. We found that FUR1-mediated uracil salvage could also rescue parasites treated with the mitochondrial complex III inhibitor atovaquone, which indirectly blocks pyrimidine synthesis by blocking ubiquinone regeneration required for *Pf*DHODH activity (**Fig. S6B**).

**Fig. 6.**
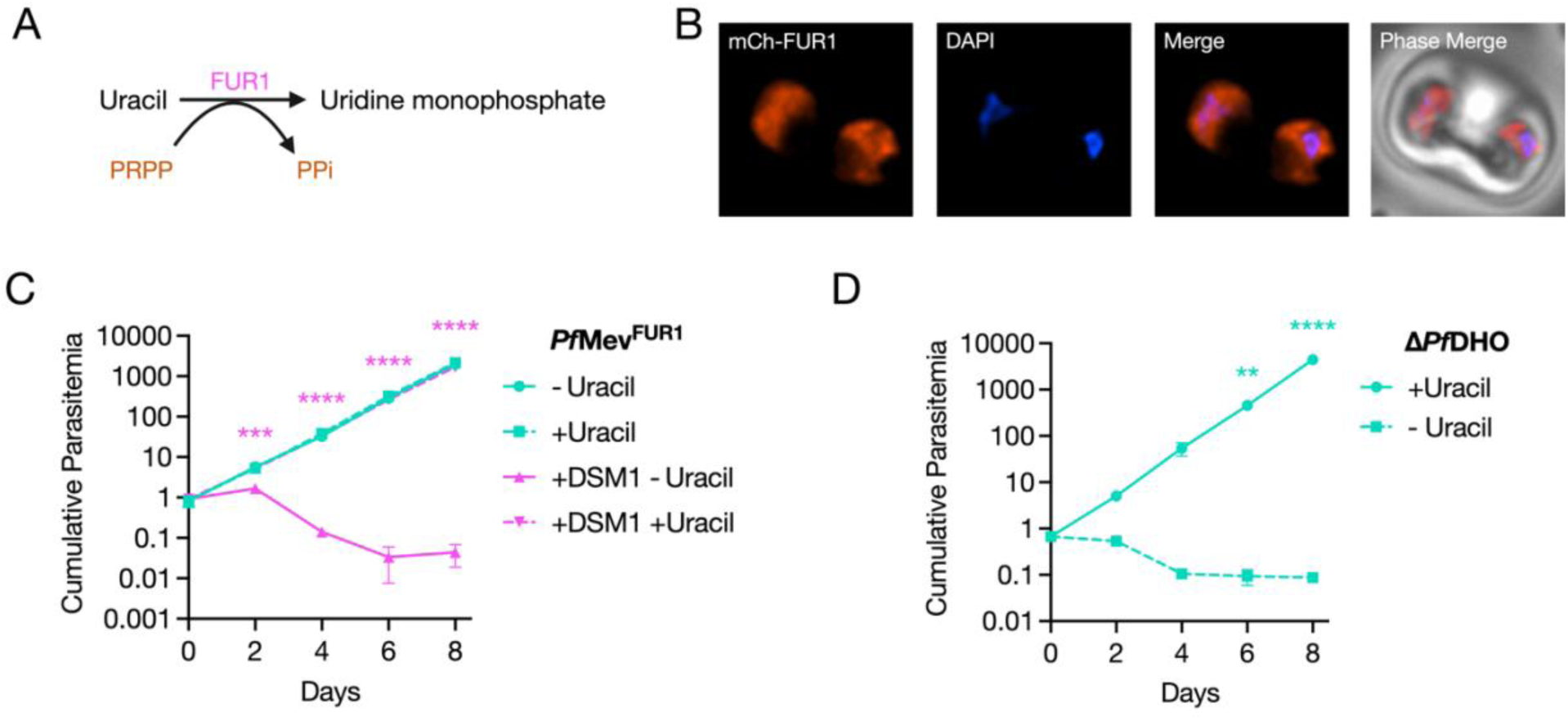
Introduction of a pyrimidine salvage enzyme in *P. falciparum* demonstrates that *Pf*DHO is essential for blood-stage growth. (A) The yeast enzyme FUR1 (uracil phosphoribosyltransferase) transfers a phosphoribosyl group from 5-phosphoribosyl-α-1-pyrophosphate (PRPP) to uracil to yield uridine 5’-monophosphate (UMP), the same product generated by the pyrimidine biosynthetic pathway. (B) Live fluorescence microscopy confirmed the expression of the mCherry-tagged FUR1 (mCh-FUR1) protein in *Pf*Mev^FUR1^ parasites. Images represent fields that are 10 μm long by 10 μm wide. (C) Growth of *Pf*Mev^FUR1^ parasites in the presence or the absence of uracil and DSM1 was measured by flow cytometry for 8 days. Growth inhibition from DSM1 treatment was averted when *Pf*Mev^FUR1^ parasites were supplemented with uracil. (D) Growth measurements of the Δ*Pf*DHO line across an 8-day period showed that the parasites relied on uracil salvage for survival. Removal of uracil from the culture medium led to rapid parasite death. Growth curves in (C) and (D) were generated from two biological experiments conducted in quadruplicate. Data were plotted on a logarithmic Y-axis. Error bars represent standard deviations from the mean. Statistical analysis was conducted using GraphPad Prism 10 (GraphPad Software, Inc). Two-way ANOVA was followed by Bonferroni’s correction (**, P ≤ 0.01; ***, P ≤ 0.001; ****, P ≤ 0.0001). Asterisks in (C) depict significant differences between +DSM1 - Uracil and +DSM1 +Uracil conditions.

In the *Pf*Mev^FUR1^ background, we successfully generated a Δ*Pf*DHO line in the presence of uracil using the same CRISPR approach that had failed in the parental *Pf*Mev line (**Fig. S7A-C**). Withdrawal of uracil led to a rapid arrest in the growth of Δ*Pf*DHO parasites, demonstrating that *Pf*DHO is indeed essential for pyrimidine synthesis (**Fig. 6D**).

### Pyrimidine biosynthetic enzymes do not have essential moonlighting functions

*Pf*ATC and *Pf*CPSII contain unusual features that raise the possibility of additional, noncanonical functions. Supporting this idea, studies in the related apicomplexan parasite *Toxoplasma gondii*, which can synthesize and salvage pyrimidines, indicate that enzymes in the pyrimidine biosynthesis pathway may serve secondary roles. Replacement of the DHODH gene in *T. gondii* with a catalytically inactive version was lethal despite intact pyrimidine salvage, pointing to a potential structural or nonenzymatic role for DHODH (54).

To test whether *Pf*ATC, *Pf*CPSII, and *Pf*DHODH serve similarly expanded roles in *P. falciparum*, we utilized the FUR1/uracil system to bypass the parasite’s reliance on its *de novo* pyrimidine synthesis pathway. The mCh-FUR1 expression plasmid was integrated into the *att*B sites in the existing *Pf*ATC^CD^ and *Pf*CPSII^CD^ parasite lines (**Fig. S8A-B**). We also established an NF54^attB^ parasite line expressing mCh-FUR1 (NF54^FUR1^) (**Fig. S9A-B**). Attempts to generate HA-tagged *Pf*DHODH conditional knockdown parasites in the NF54^FUR1^ background were unsuccessful. Since peptide tags can affect the structure and function of some proteins, we repeated the transfections using a TetR-DOZI plasmid modified to omit the addition of the C-terminal HA tag to *Pf*DHODH. This approach successfully yielded a *Pf*DHODH^CD^ parasite line (**Fig. S9C-D**). All three conditional knockdown lines were maintained in media containing aTet.

If *Pf*ATC, *Pf*DHO, and *Pf*DHODH play essential roles beyond pyrimidine synthesis, we would expect that the FUR1/uracil salvage mechanism would not compensate for their loss. To test this, parasites from the three transgenic lines were divided into media with or without aTet and uracil. Parasite growth was monitored over an 8-day period. As expected, all three lines failed to grow in the absence of aTet and uracil (**Fig. 7A-C**). However, the presence of uracil in aTet-deprived cultures completely restored the growth of the *Pf*ATC^CD^, *Pf*CPSII^CD^, and *Pf*DHODH^CD^ parasites, indicating that the essential function of these enzymes is confined to pyrimidine synthesis during asexual blood-stage replication.

**Fig. 7.**
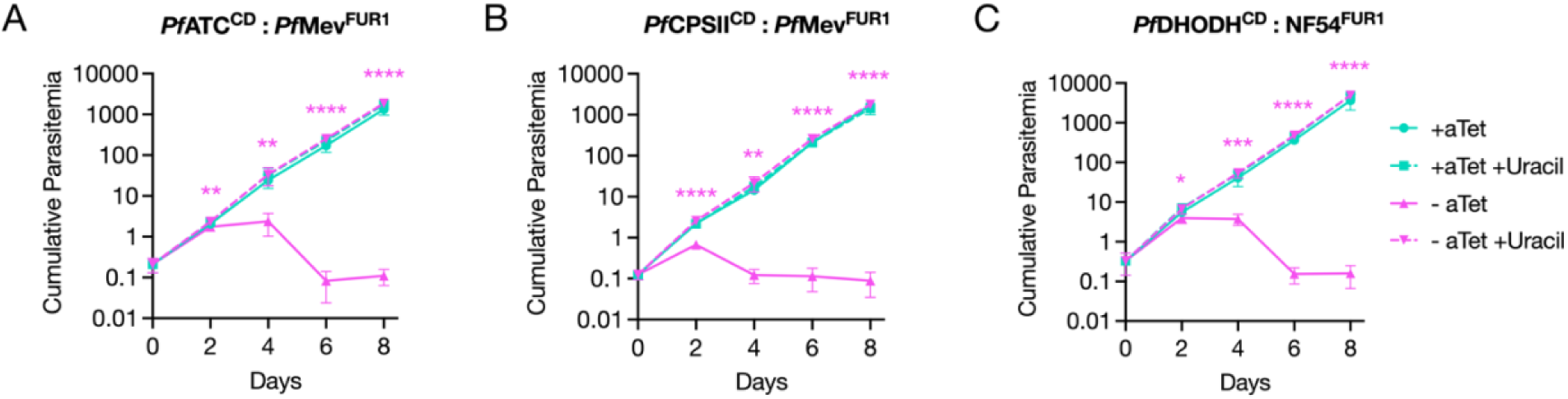
*Pf*ATC, *Pf*CPSII and *Pf*DHODH are dispensable in the presence of a pyrimidine salvage pathway. (A) *Pf*ATC^CD^, (B) *Pf*CPSII^CD^, and (C) *Pf*DHODH^CD^ parasites expressing mCh-FUR1 were monitored for their growth kinetics in the presence or absence of aTet and uracil by flow cytometry for 8 days. Parasite death resulting from the removal of aTet could be prevented with uracil supplementation. Growth curves were generated from two biological experiments were conducted in quadruplicate. Error bars represent standard deviations from the mean. Two-way ANOVA was performed using GraphPad Prism 10 (GraphPad Software, Inc) followed by Bonferroni’s correction, with significant differences noted between - aTet and – aTet +Uracil conditions. (*, P ≤ 0.05; **, P ≤ 0.01; ***, P ≤ 0.001; ****, P ≤ 0.0001).

Finally, we used the *Pf*ATC^CD^:*Pf*Mev^FUR1^ parasites generated for this experiment to validate the specificity of the α-*Pf*ATC antibody described earlier (**Fig. 2**), taking advantage of the fact that these parasites remain viable with uracil supplementation even after *Pf*ATC knockdown. Immunofluorescence microscopy *Pf*ATC signal was detected in cultures maintained with aTet and uracil, but this signal was largely lost upon aTet withdrawal, confirming both antibody specificity and effective protein depletion (**Fig. S10**).

## DISCUSSION

In this report, we applied molecular genetic approaches to study multiple enzymes in the pyrimidine biosynthetic pathway of *Plasmodium falciparum*. This pathway, which contains the established drug target *Pf*DHODH, exhibits significant structural differences from that of the mammalian host. In humans, the first three and last two reactions are catalyzed by the multifunctional proteins CAD and UMP synthase, respectively (34, 35). By contrast, *P. falciparum* encodes all six pathway enzymes as discrete gene products, with some containing unknown features that could indicate additional functions or distinct modes of regulation (11, 14, 33, 37).

To determine the importance of these parasite-specific differences, we initially focused our analysis on *Pf*ATC, which has been structurally resolved and characterized *in vitro* (19, 65). Using a conditional knockdown approach, we confirmed that *Pf*ATC is essential for parasite survival (**Fig. 3**). Contrary to predictions that its N-terminal extension functions as an apicoplast-trafficking peptide, antibodies raised against *Pf*ATC showed that the protein localizes primarily to the cytoplasm (**Fig. 2D-E**). Interestingly, both the N-terminal peptide alone and full-length *Pf*ATC expressed as a second copy from a dispensable locus were able to direct a fluorescent reporter to the parasite periphery, suggesting that low-level or stage-specific trafficking to this compartment may occur (**Fig. 2A-B**). We have not examined if peripheral *Pf*ATC associates with the parasite plasma membrane or is present in the parasitophorous vacuole. This observed discrepancy in localization between native *Pf*ATC and the tagged transgenes could reflect variations in expression level or timing arising from the use of a heterologous promoter. Deletion of the N-terminus from the second copy abolished peripheral localization and diminished its ability to rescue parasites depleted of endogenous *Pf*ATC (**Fig. 4**). This indicates that the N-terminal region contributes to *Pf*ATC function, potentially by influencing protein targeting, stability, or interactions with other proteins or membranes. If some endogenous *Pf*ATC indeed localizes to the parasite periphery, a plausible model is that its N-terminal region acts as a membrane anchor to organize *Pf*CPSII and *Pf*DHO into a higher-order complex analogous to the multifunctional architecture of human CAD, thereby enhancing metabolic efficiency (34). Notably, a recent spatial proteomic map of *P. falciparum* schizonts assigned *Pf*ATC to the plasma membrane, lending circumstantial support to this model (66).

*Pf*ATC has been the focus of several drug discovery efforts (65, 67–70). Structural analyses of the apo (T-state) and liganded (R-state) forms of the enzyme laid the groundwork for rational drug design aimed at disrupting the transition between the two states (65). More recent fragment-based studies have revealed a previously unrecognized allosteric pocket in *Pf*ATC (68). A series of compounds selectively binding to this site were found to block parasite growth in red blood cells with limited effects against human cells (68). These advances, together with our genetic validation of *Pf*ATC essentiality, strengthen the rationale for targeting this enzyme for antimalarial drug development.

*Pf*CPSII, which catalyzes the first reaction in the pathway, is among the largest known members of this enzyme class due to two insertions of unknown function (14). As predicted, *Pf*CPSII was found to be essential for parasite growth (**Fig. 5D**). C-terminal epitope tagging revealed that *Pf*CPSII is cytoplasmic and produces multiple bands on an immunoblot, indicative of protein processing or degradation (**Fig. 5C**). Proteolytic processing may be a necessary step in CPSII maturation, generating different enzymatic or regulatory subunits from the large precursor protein. Future work involving protein truncation analysis will be important to define the functional contributions of the insertions.

*Pf*DHO was previously predicted to be dispensable, albeit with a fitness cost, in a genome-wide transposon mutant screen (41). One possible explanation for this phenotype is that carbamoyl aspartate undergoes slow, spontaneous hydrolysis to form dihydroorotate in the absence of *Pf*DHO, thereby permitting limited growth. To directly test the essentiality of *Pf*DHO, we bypassed the parasite’s reliance on *de novo* pyrimidine biosynthesis by introducing the yeast uracil phosphoribosyltransferase FUR1, which converts uracil to UMP (**Fig. 6A**) (64). This system allowed us to modulate the activity of the salvage enzyme by adjusting uracil concentrations in the culture medium. As proof of concept, we demonstrated that uracil supplementation rescued parasites from inhibition by DSM1 or atovaquone (**Fig. 6C**, **Fig. S6B**). Using this approach, we showed that *Pf*DHO is essential, as knockout parasites survived only when supplied with exogenous uracil (**Fig. 6D**). Functionally, FUR1 achieves the same outcome as yeast DHODH (yDHODH) in circumventing ubiquinone-dependent pyrimidine synthesis, but its uracil-driven activity provides a controllable alternative to the constitutive bypass from yDHODH. This makes the FUR1 system a versatile genetic tool, both as a selectable marker and for screening compounds that target the mETC and pyrimidine biosynthetic enzymes.

We next applied this system to investigate whether the additional domains in *Pf*ATC and *Pf*CPSII confer functions beyond their canonical enzymatic roles. By introducing FUR1 into conditional knockdown lines and regulating uracil availability, we could selectively rescue pyrimidine synthesis and assess whether these enzymes contribute to parasite fitness through other mechanisms. In both cases, uracil supplementation fully restored growth, indicating that *Pf*ATC and *Pf*CPSII are essential at this stage of parasite development solely for their roles in *de novo* pyrimidine biosynthesis (**Fig. 7A-B**). We extended this approach to determine if *Pf*DHODH, like its homolog in *T. gondii* (54), also possesses a non-enzymatic role. In *P. falciparum*, however, *Pf*DHODH appears to function exclusively in pyrimidine synthesis during parasite replication in red blood cells (**Fig. 7C**).

Overall, our findings strengthen the case for prioritizing the pyrimidine biosynthesis pathway in antimalarial drug discovery. The distinct domain architecture of its constituent enzymes makes them attractive for the development of highly specific inhibitors. Particularly, it will be of interest to further study *Pf*CPSII to determine if its unusual insertions confer distinct mechanisms of regulation or maturation that could be exploited therapeutically.

## METHODS

### Parasite culture

*P. falciparum* NF54^attB^ and *Pf*Mev^attB^ parasites were cultured in red blood cells (RBCs) at 2% hematocrit using complete medium supplemented with AlbuMAX II (CMA) (58, 59). CMA was prepared from RPMI-1640 medium with L-glutamine (USBiological Life Sciences, R8999) and further supplemented with 0.2% sodium bicarbonate, 12.5 μg/mL hypoxanthine, 20 mM HEPES, 5 g/L AlbuMAX II (ThermoFisher Scientific, 11021037), and 25 μg/mL gentamicin. Cultures were maintained in flasks or plates at 37°C under a gas mixture of 94% N₂, 3% O₂, and 3% CO₂.

### Plasmid construction

Various *Pf*ATC-mCherry expression vectors were generated using the previously described p15-*Ec*DPCK-mCherry plasmid as a backbone (71). To replace the *Ec*DPCK gene, the plasmid was digested with *Avr*II and *Bsi*WI. Coding sequences for *Pf*ATC_36_, *Pf*ATC_375_, or *Pf*ATC_Δ36_ were amplified from *P. falciparum Pf*Mev^attB^ cDNA using primers Pyr_001 - Pyr_004 containing flanking *Avr*II and *Bsi*WI sites (**Table S1**). The PCR products were then digested with the corresponding enzymes and ligated into the linearized p15-mCherry vector.

The backbone from plasmid p15-*Ec*DPCK-mCherry (71) was also used to generate the p15-mCherry-FUR1 plasmid by replacing the region encoding *Ec*DPCK-mCherry with nucleotides encoding mCherry-FUR1. The mCherry fragment was amplified from the p15-*Pf*ATC_36_-mCherry vector using primers Pyr_005 and Pyr_006 (**Table S1**), containing *Avr*II sites, and the *S. cerevisiae* uracil phosphoribosyltransferase gene FUR1 was synthesized by LifeSct LLC (Rockville, MD) and amplified using primers (Pyr_007 and Pyr_008) containing *Bsr*GI and *Xho*I restriction sites (**Table S1**).

To knock down gene expression, we used the pKD plasmid and its derivatives, which contain the TetR-DOZI expression cassette and a 10x aptamer array (61). Homology arms 1 (HA1) and 2 (HA2) for *Pf*ATC, *Pf*CPSII, and *Pf*DHODH were amplified using primers listed in **Table S1**. The reverse primers were designed to contain a recodonized sequence to eliminate homology with the guide RNA. Additional HA1 reverse primers were used for *Pf*ATC and *Pf*CPSII to restore the native coding sequence downstream of the recodonized region, but not including the stop codon. HA2 amplicons were cloned into the *Asc*I and *Eco*RV sites in pKD, and the final HA1 amplicons were ligated into *Eco*RV and *Aat*II sites. For *Pf*DHO, a synthetic fragment consisting of HA2-recoded HA1-3×FLAG flanked by *Asc*I and *Psp*OM1 sites (LifeSct LLC) was cloned into the pTDN plasmid, a derivative of pKD (**File S2**, **Table S1)**. pTDN has three changes relative to pKD. Unnecessary attP sites were removed by digestion with *Eco*RI, followed by religation of the plasmid. The TetR-DOZI coding sequence was replaced with a synthesized sequence (LifeSct LLC, **File S2**) to remove unwanted endonuclease sites and inserted into the *Avr*II and *Nhe*I sites in pKD. Finally, the blasticidin deaminase gene was replaced with a synthetic neomycin resistance gene (LifeSct LLC, **File S2**) via digestion with *Age*I and *Ngo*MIV, followed by ligation. To create a version of the *Pf*DHODH-pKD plasmid lacking the C-terminal 2×FLAG tag, the HA1 sequence along with the stop codon was amplified using primers encoding *Eco*RV and *Psp*OM1 sites (**Table S1**), digested with the corresponding enzymes, and cloned into the same sites in the *Pf*DHODH-pKD vector, effectively removing the tag.

The gene deletion construct for *Pf*DHO was generated using the pRSng-BD repair plasmid (72). The HA1 and HA2 regions were amplified using primers listed in **Table S1**. The HA1 sequence was inserted into the *Not*I site and HA2 into the *Ngo*MIV site of the plasmid.

To construct plasmids encoding guide RNAs for gene knockdown or knockout experiments, the pCasG-BD-LacZ vector was first digested with the restriction enzyme *Bsa*I (61). Custom guide RNA sequences were synthesized as complementary oligonucleotides (**Table S1**), annealed to form duplexes, and subsequently cloned into the digested vector by ligation-independent cloning with In-Fusion (Takara, 639650).

All restriction enzymes and T4 DNA Ligase (M0202S) were sourced from New England Biolabs.

### Parasite transfections

To generate transgenic *P. falciparum* lines expressing *Pf*ATC-mCherry variants and mCherry-FUR1, 350 μL volumes of RBCs were electroporated using the GenePulser XCell system (Bio-Rad) with: 50 μg of the pINT plasmid (59) that expresses the Bxb1 integrase, and 65 μg of the p15 plasmid with an expression cassette for the mCherry fusion gene and the dihydrofolate reductase (DHFR) resistance marker that confers resistance to the compound WR99210. Electroporated RBCs were mixed with 1 mL of parasite culture at 2-3% parasitemia and maintained in 10 mL of CMA for 2 days, then transferred to CMA containing 2.5 nM WR99210 (Jacobus Pharmaceutical Company, Inc.) for 7 days. Cultures were then returned to drug-free CMA until parasites became detectable by blood smear, at which point WR99210 was reintroduced.

For knockdown lines, RBCs were electroporated with 65 μg each of the pKD/pTDN plasmid and the corresponding pCasG-BD-gRNA plasmid. Following electroporation, the RBCs were treated as above and maintained in CMA with 0.5 μM anhydrotetracycline (aTet; Cayman Chemical: 10009542). Cultures were placed under selection with either 2.5 μg/mL blasticidin (for pKD; Corning: 30-100-RB) or 0.5 mg/mL G418 (for pTDN; Corning: 61-234-RG) along with aTet for 7 days. They were then maintained in CMA and aTet until parasites reemerged, after which drug selection was resumed.

For *Pf*DHO gene deletion, RBCs were electroporated with the pRSngBD-*Pf*DHO plasmid and the corresponding pCasG-gRNA plasmid, then mixed with 1 mL of *Pf*Mev^FUR1^ parasites at 2-3% parasitemia in CMA supplemented with 50 μM uracil. Two days post transfection, the culture was selected with 2.5 μg/mL blasticidin in the presence of uracil for 7 days, followed by maintenance in CMA and uracil until parasite reappearance, at which point blasticidin was added back to the medium.

Clonal parasites for all transgenic lines were obtained by limiting dilution in 96-well plates. The genotypes of transgenic parasites were confirmed by PCR using primers listed in **Table S1**.

### Growth assays

To assess the growth kinetics of various parasite lines under different treatments, the parasites were washed three times to remove existing drugs and supplements from the media, diluted to 0.5% parasitemia in 2% hematocrit, and distributed into the appropriate selective media conditions. Cultures were seeded in 200 μL volumes per well in 96-well plates (ThermoFisher Scientific, 267427), and the parasitemia was determined by flow cytometry every other day over an 8-day period, unless stated otherwise. Immediately after each sampling point, cultures diluted eight-fold to prevent overgrowth. For the data represented in **Fig. 5D** and **Fig. S6B**, cultures were instead diluted four-fold and ten-fold every two days, respectively. All source data for the growth curves are available in **File S3**.

To prepare samples for flow cytometry, 20 μL volumes of parasites were taken from each well, diluted 1:5 in PBS and stored at 4 °C in 96-well plates until analysis. Parasites were stained by adding 10 μL of the 1:5 dilutions to a 96-well plate containing 100 μL of 1×SYBR Green (Invitrogen, S7563) in PBS, followed by a 30-minute incubation in the dark on a plate shaker. Afterwards, 150 μL of PBS was added to reduce background fluorescence from unbound dye. Stained cells were analyzed using an Attune NxT Flow Cytometer (ThermoFisher Scientific), operated at a flow rate of 50 μL/minute to collect 10,000 events per sample. Each condition was tested in quadruplicate.

### Antibody generation

DNA encoding amino acid residues 38–375 of *Pf*ATC was PCR-amplified from *P. falciparum* NF54^attB^ genomic DNA using primers Pyr_064 and Pyr_065 (**Table S1**). The product was digested with *Eco*RI and *Hin*dIII and cloned into the same sites of the pMALcHT *Escherichia coli* expression vector (73) using In-Fusion ligation-independent cloning to yield the pMALcHT-*Pf*ATC expression plasmid. This plasmid was cotransformed with the pRIL plasmid purified from BL21-CodonPlus (DE3)-RIL competent cells (Agilent Technologies), and plasmid pKM586 encoding TEV protease (74). When coexpressed, the TEV protease allows for *in vivo* cleavage of the N-terminal maltose binding protein (MPB) from the MBP-6xHis-*Pf*ATC.

*E. coli* cells transformed with the plasmids above were grown in LB medium at 37 °C to an OD_600_ of 0.6, at which point protein expression was induced with 0.4 mM isopropyl β-thiogalactopyranoside (IPTG). Cells were harvested after 8 hours of growth at 16 °C. Cell pellets were resuspended in 20 mL of lysis buffer (10 mM Tris-HCl, pH 8.0, 100 mM NaCl, 2.5 µg/mL DNase I, and 1 mg/mL lysozyme) per liter of cell culture. The resuspended cells were lysed by sonication, and the lysate was clarified by centrifugation at 4 °C for 20 minutes at 20,000 × g. The supernatant was loaded onto a Ni^2+^-charged 5 mL HiTrap chelating HP column (Cytiva) pre-washed with equilibration buffer (10 mM Tris-HCl, pH 8.0, 100 mM NaCl). The column was washed with 2 column volumes of equilibration buffer before connecting to a 5 mL MBPTrap HP column (Cytiva) pre-washed with equilibration buffer (the MBPTrap column will remove any uncut fusion protein). The bound proteins from the HiTrap column were eluted using a 0 - 400 mM imidazole gradient in equilibration buffer, and further purified by size-exclusion chromatography using a HiPrep 26/60 Sephacryl S-100 HR size-exclusion column (Cytiva) equilibrated with phosphate-buffered saline (PBS, pH 7.4).

Pure recombinant *Pf*ATC was used to generate rat antiserum (Cocalico Biologicals Inc.) Two rats were immunized with *Pf*ATC in Complete Freund’s Adjuvant and were boosted at weeks 2, 3, and 7. Terminal bleeds were collected on day 56. Anti-*Pf*ATC IgG was purified from antiserum using a *Pf*ATC affinity column as described previously (75). A total of 10 mg of α-*Pf*ATC IgG was concentrated to 0.25 mg/mL and stored at −80 °C in storage buffer (PBS, 40% glycerol, 0.02% NaN₃).

### Western Blotting

Parasite pellets were collected by centrifugation at 500 × g for 5 minutes at room temperature (RT). To release parasites from RBCs, the pellets were treated with 0.15% saponin for 5 minutes at RT, followed by three washes in 1×PBS. The resulting parasite pellets were solubilized in NuPAGE LDS sample buffer (Invitrogen: NP0007) supplemented with 10% β-mercaptoethanol and heated at 95 °C for 5 minutes. Protein extracts were separated on 4–12% Bis-Tris SDS-PAGE gels (Invitrogen: NP0322) and transferred onto nitrocellulose membranes. Membranes were blocked with 5% non-fat milk in 1×PBST (PBS with 0.1% Tween20) and incubated overnight at 4 °C with the appropriate primary antibodies at the following dilutions in milk/PBST: 1:20,000 mouse α-FLAG M2 monoclonal antibody (Millipore Sigma: F3165) 1:2,500 rat α-HA monoclonal antibody (3F10, Roche: 11867423001), 1:250 affinity purified rat α-*Pf*ATC polyclonal antibodies (this study), 1:500 rat α-*Pf*ATC antisera, or 1:1,000 rabbit α-*Pf*BiP antisera (BEI Resources: MRA-1246) (76, 77). After washing, blots were incubated at room temperature for 1 hour with HRP-conjugated secondary antibodies from GE Healthcare (1:10,000 anti-mouse, 1:5000 anti-rat, or 1:5000 anti-rabbit). Signal was developed using SuperSignal West Pico Chemiluminescent Substrate (ThermoFisher Scientific: 34580) and visualized on X-ray film. To assess loading controls, blots were stripped using 200 mM glycine (pH 2.0) for 5 minutes, then reprobed overnight at 4 °C.

### Microscopy

For live fluorescence microscopy, 100 μL of parasite culture was stained with 1 μg/mL DAPI (ThermoFisher Scientific: 62248) for 30 minutes at 37 °C, washed three times with CMA, and resuspended in 10 μL CMA for mounting on glass slides. Coverslips were sealed with wax, and images spanning 5 μm (0.2 μm steps) were acquired using a Zeiss AxioImager M2 microscope. For immunofluorescence assays, parasites were fixed in 4% formaldehyde and 0.0075% glutaraldehyde in PBS for 30 minutes, permeabilized with 1% Triton X-100, and blocked in 3% BSA for 2 hours. Cells were incubated overnight at 4 °C with 1:200 affinity purified rat α-*Pf*ATC antibodies (this study), 1:500 rabbit α-ACP antibodies (78), 1:500 rabbit α-BiP antibody (BEI Resources: MRA-1246), or 1:500 mouse α-FLAG M2 antibody (Millipore Sigma: F3165), followed by PBS washes and incubation with Alexa Fluor-conjugated secondary antibodies: 1:300 goat anti-rat 488 (ThermoFisher Scientific: A-11006), 1:500 goat anti-rabbit 594 (ThermoFisher Scientific: A-11037), or 1:500 goat anti-mouse 488 (ThermoFisher Scientific: A-11001). Samples were mounted in ProLong™ Gold Antifade reagent with DAPI (Invitrogen: P36935) and sealed with nail polish. For both live and fixed imaging, Z-stacks were deconvolved using VOLOCITY software (PerkinElmer), and representative single-plane images were reported.

## ACKNOWLEDGEMENTS

This work was supported by NIH R01 AI125534 (awarded to S.T.P.), the Johns Hopkins Malaria Research Institute (JHMRI), and the Bloomberg Family Foundation. K.R. was supported by the Johns Hopkins Discovery Award. M.L.S. was supported by NIH training grant T32AI007417. R.E. was supported by the JHMRI postdoctoral fellowship and the Samuel Jordan Graham postdoctoral fellowship. We acknowledge VEuPathDB for providing critical data resources that supported this study.

