## Supplementary material for "Genetic analysis of pyrimidine biosynthetic enzymes in *Plasmodium falciparum*": File S1

A

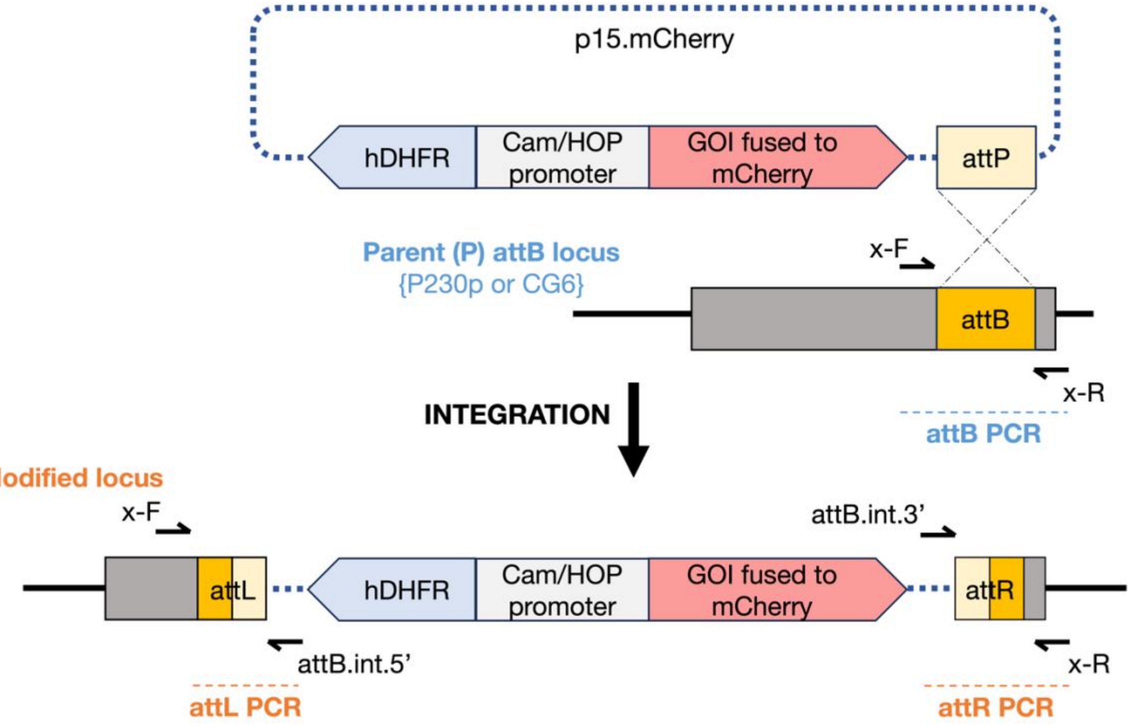


B

| Lane | PCR reaction | Primer Combination x = P230p | | Product size (bp) | Parasite Line |
|---|---|---|---|---|---|
| 1 | attL | x-F Pyr_013 | attB.int.5' Pyr_011 | 537 | Modified |
| 2 | attR | attB.int.3' Pyr_012 | x-R Pyr_014 | 487 | |
| 3 | attB | x-F Pyr_013 | x-R Pyr_014 | – | |
| 4 | attL | x-F Pyr_013 | attB.int.5' Pyr_011 | – | Parent (P) |
| 5 | attR | attB.int.3' Pyr_012 | x-R Pyr_014 | – | |
| 6 | attB | x-F Pyr_013 | x-R Pyr_014 | 1241 | |

C

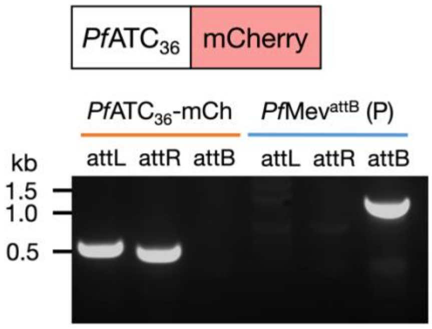


D

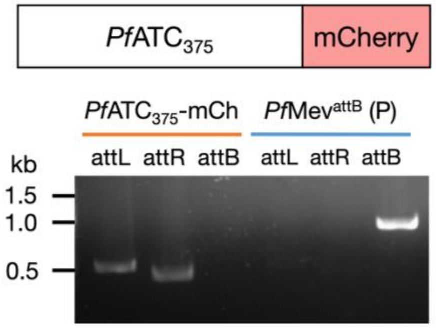

**Fig. S1. Generation of transgenic P*. falciparum* lines with mCherry-tagged proteins of interest.** (A) The schematic depicts integration of the p15.mCherry plasmid into the parasite genome. The p15.mCherry plasmid was inserted into the P230p or CG6 locus of *P. falciparum Pf*Mev$^{attB}$ or NF54$^{attB}$ parasites, respectively, by Bxb1 integrase-mediated recombination at the plasmid *att*P and genome *att*B sites. The modified locus contains the entire plasmid flanked by new *att*L and *att*R sites that are generated by recombination. Arrows indicate the positions of primers used for diagnostic PCR. The Cam/HOP bidirectional promoter drives expression of the gene of interest (GOI) fused to mCherry and the selection marker hDHFR (human dihydrofolate reductase). (B) Primer pairs for verifying integration of the p15.mCherry plasmid into the P230p genomic locus are listed along with the expected sizes of the PCR products. Integration of (C) p15.*Pf*ATC$_{36}$-mCherry, and (D) p15.*Pf*ATC$_{375}$-mCherry plasmids in *Pf*Mev$^{attB}$ (Parent, 'P') parasites was confirmed by PCR amplification (attL and attR products). The intact P230p locus (attB product) was detected only in the parent (blue) and not in the transgenic lines (orange).

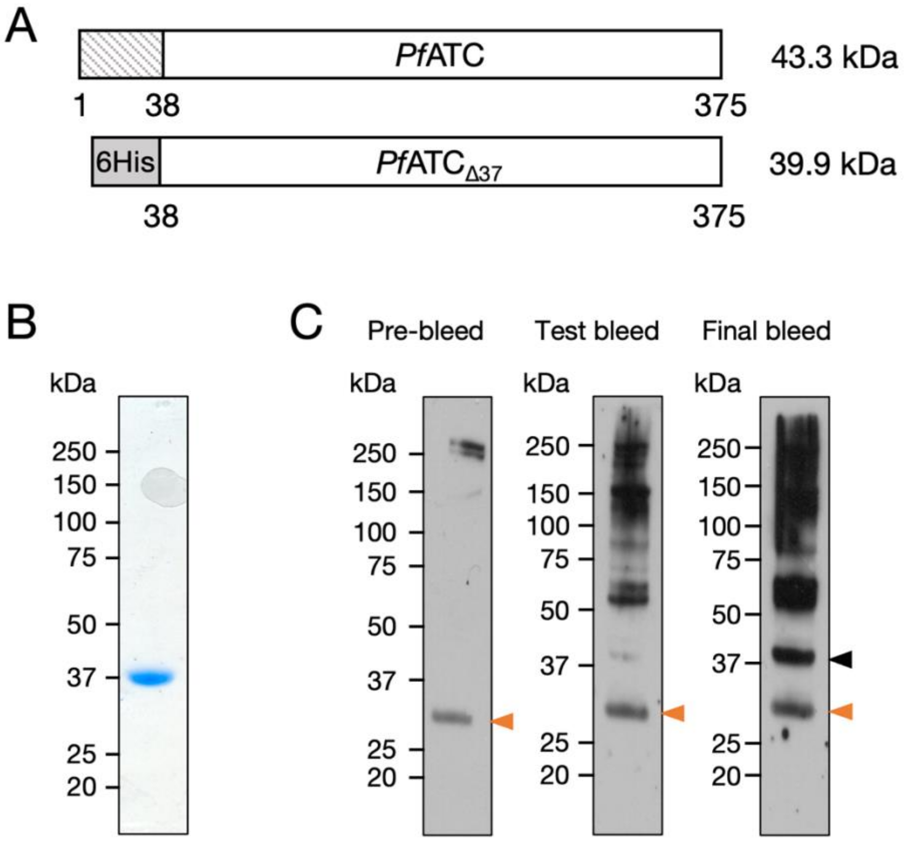


**Fig. S2. Generation and validation of *Pf*ATC antisera.** (A) *Pf*ATC (amino acids 38–375) was fused to an N-terminal 6×His tag for recombinant protein expression and purification. (B) Purified recombinant *Pf*ATC resolved on SDS-PAGE under reducing conditions and stained with Coomassie blue. (C) Purified *Pf*ATC was probed with pre-bleed (day 0), test bleed (day 35), or final bleed (day 56) antisera (1:500) from a rat immunized with recombinant *Pf*ATC. The black arrowhead indicates the specific band for *Pf*ATC, while the orange arrowhead marks a non-specific band detected even with pre-bleed antisera.

A

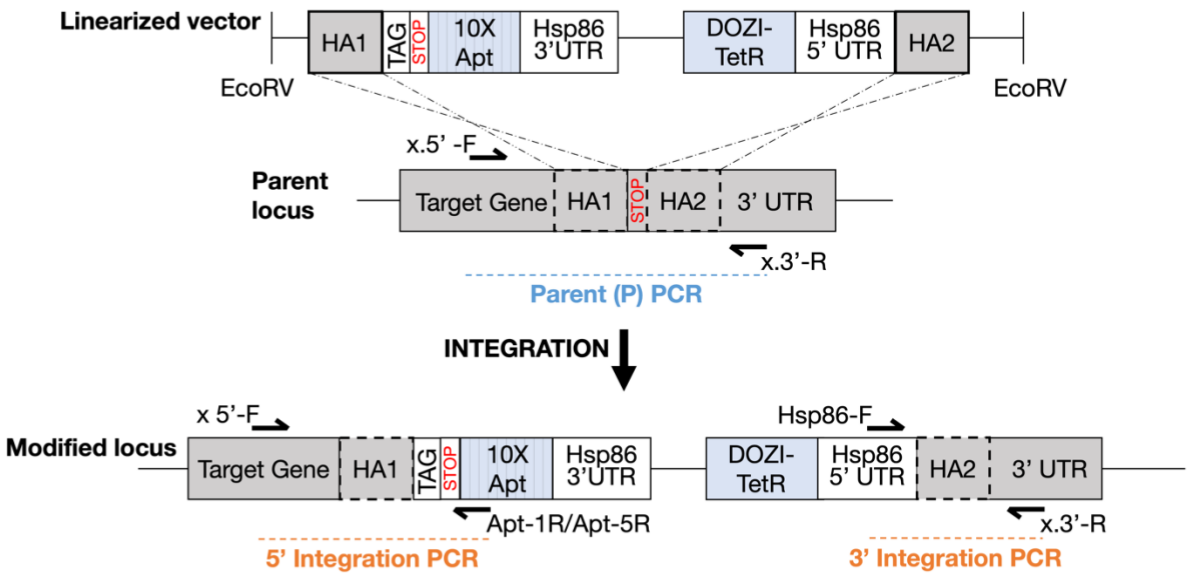


B

| Lane | PCR reaction | Primer Combination x = *Pf*ATC | | Product size (bp) | Parasite Line |
|---|---|---|---|---|---|
| 1 | 5' | x.5'-F Pyr_040 | Apt-1R Pyr_048 | 896 | *Pf*ATC$^{CD}$ |
| 2 | 3' | Hsp86-F Pyr_049 | x.3'-R Pyr_041 | 766 | |
| 3 | P | x.5'-F Pyr_040 | x.3'-R Pyr_041 | – | |
| 4 | 5' | x.5'-F Pyr_040 | Apt-1R Pyr_048 | – | *Pf*Mev$^{attB}$ Parent (P) |
| 5 | 3' | Hsp86-F Pyr_049 | x.3'-R Pyr_041 | – | |
| 6 | P | x.5'-F Pyr_040 | x.3'-R Pyr_041 | 1610 | |

C

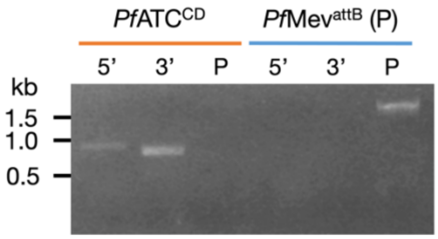


**Fig. S3. Schematic of knockdown plasmid insertion into the 3' UTR of target genes.** (A) The pKD plasmid was linearized by digestion with *Eco*RV prior to transfection. Homologous regions between the linearized vector and the Parent (P) locus are indicated by dotted lines. The Hsp86 5' UTR (untranslated region)

contains a promoter element that drives TetR-DOZI expression. Arrows indicate the positions of primers used for diagnostic PCR. 'TAG' refers to the 2×FLAG epitope tag appended to the target protein, and 10×Apt refers to the aptamer array inserted in the 3' UTR of the target gene. HA1 and HA2 are the homology arms utilized for recombination. This schematic also applies to the pTDN plasmid. (B) Primer pairs for verifying integration of the pKD-*Pf*ATC plasmid into the 3' UTR of the *Pf*ATC gene are listed, along with the expected sizes of the PCR products. (C) Integration of the pKD-*Pf*ATC plasmid was confirmed by PCR amplification of the 5' and 3' loci at the insertion site in *Pf*ATC$^{CD}$ clonal parasites (orange). The *Pf*Mev$^{attB}$ parent line served as a control (blue).

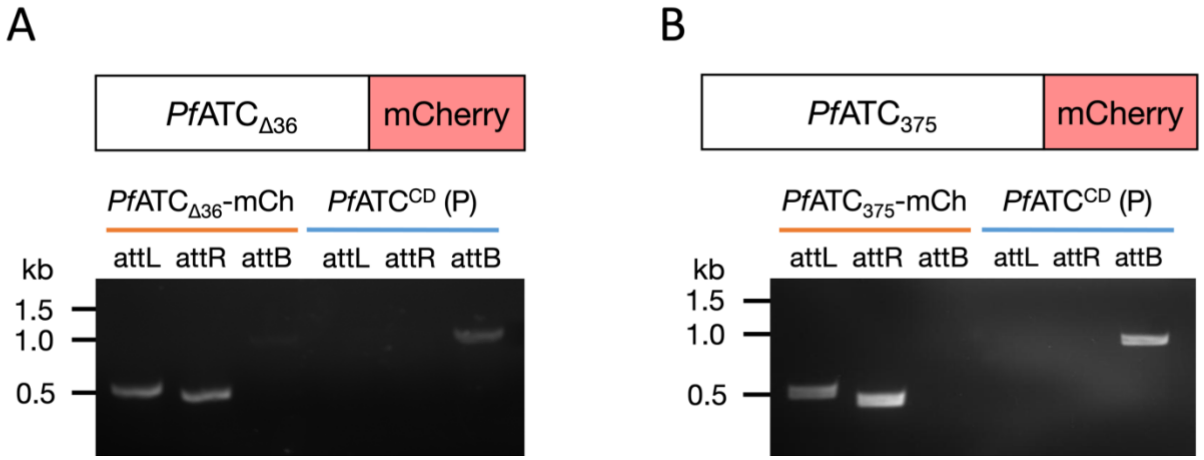


**Fig. S4. Bxb1-mediated integration of *Pf*ATC-mCherry expression constructs into the *Pf*ATC$^{CD}$ parasite genome.** Genomic integration of (A) p15.*Pf*ATC$_{\Delta 36}$-mCherry and (B) p15.*Pf*ATC$_{375}$-mCherry plasmids in *Pf*ATC$^{CD}$ (Parent, 'P') parasites was confirmed by PCR amplification (*att*L and *att*R products). The intact P230p locus (*att*B product) was detected only in the parent (blue) and not in the transgenic lines (orange). Refer to the schematic in **Fig. S1A** for a detailed illustration of the integration of p15.mCherry plasmids into *att*B sites in the *P. falciparum* genome. Primer pairs for verifying integration of the p15.mCherry plasmid into the P230p locus are listed in **Fig. S1B** along with the expected sizes of the PCR products.

A

| Lane | PCR reaction | Primer Combination | | Product size (bp) x = *Pf*CPSII | Product size (bp) x = *Pf*DHO | Parasite line |
|---|---|---|---|---|---|---|
| 1 | 5' | x.5'-F Pyr_042/Pyr_044 | Apt-1R Pyr_048 | 1074 | 1025 | x$^{CD}$ |
| 2 | 3' | Hsp86-F Pyr_049 | x.3'-R Pyr_043/Pyr_045 | 843 | 433 | |
| 3 | P | x.5'-F Pyr_042/Pyr_044 | x.3'-R Pyr_043/Pyr_045 | – | – | |
| 4 | 5' | x.5'-F Pyr_042/Pyr_044 | Apt-1R Pyr_048 | – | – | *Pf*Mev$^{attB}$ Parent (P) |
| 5 | 3' | Hsp86-F Pyr_049 | x.3'-R Pyr_043/Pyr_045 | – | – | |
| 6 | P | x.5'-F Pyr_042/Pyr_044 | x.3'-R Pyr_043/Pyr_045 | 1469 | 1014 | |

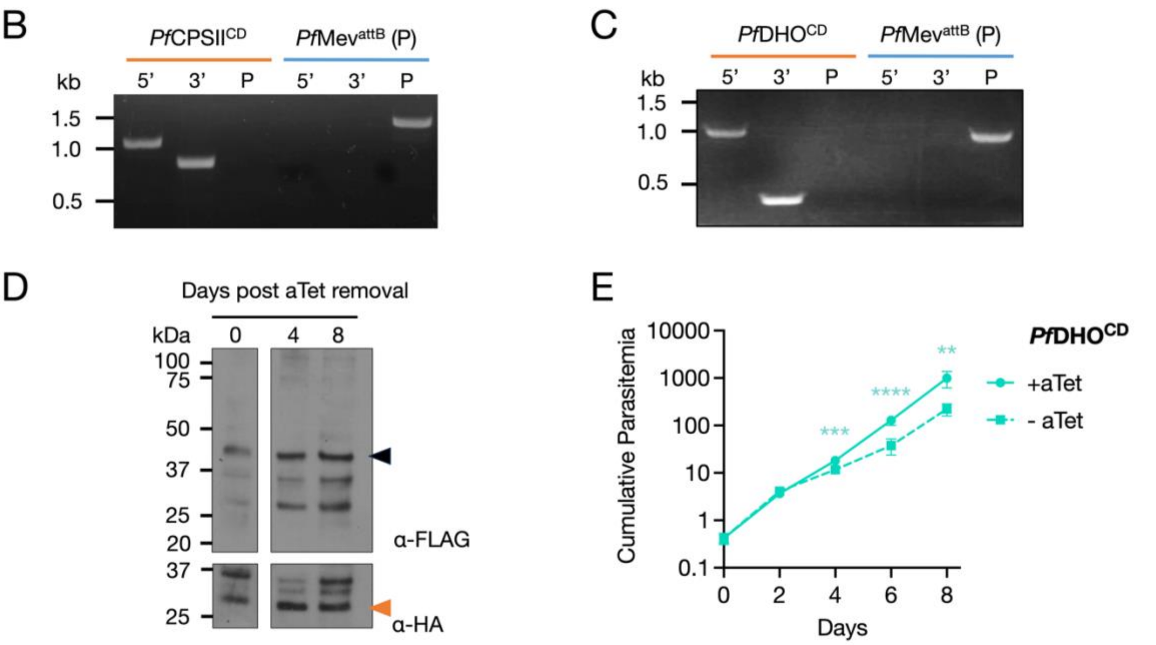


**Fig. S5. Genotyping and characterization of *Pf*CPSII and *Pf*DHO conditional knockdown lines.** (A) Primer pairs for verifying integration of the pKD-*Pf*CPSII plasmid and pTDN-*Pf*DHO plasmid into the 3' UTR of *Pf*CPSII and *Pf*DHO genes, respectively, are listed along with the expected sizes of the PCR products. (B) Integration of the pKD-*Pf*CPSII plasmid, and (C) the pTDN-*Pf*DHO plasmid to create *Pf*CPSII$^{CD}$ and *Pf*DHO$^{CD}$ parasite lines, respectively, was validated by PCR amplification of the 5' and 3' loci at the genomic insertion sites (orange). The *Pf*Mev$^{attB}$ parent (P) line served as a control (blue). Refer to **Fig. S3A** for a schematic of knockdown plasmid insertion into the 3' UTR of target genes. (D) Western blot analysis of *Pf*DHO$^{CD}$ parasites at Day 0, 4 and 8 of aTet withdrawal, using α-FLAG antibody, showed no measurable knockdown of FLAG-tagged *Pf*DHO (black arrowhead). The α-HA antibody detecting HA-tagged api-SFG (orange arrowhead) was used as a loading control. (E) Growth of *Pf*DHO$^{CD}$ parasites in the presence or the absence of aTet was monitored by flow cytometry for 8 days. Removal of aTet led to a reduction in parasite growth. Two biological experiments were conducted in quadruplicate (two-way ANOVA computed using GraphPad Prism 10, followed by Bonferroni's correction; **, P ≤ 0.01; ***, P ≤ 0.001; ****, P ≤ 0.0001). Error bars indicate standard deviations from means.

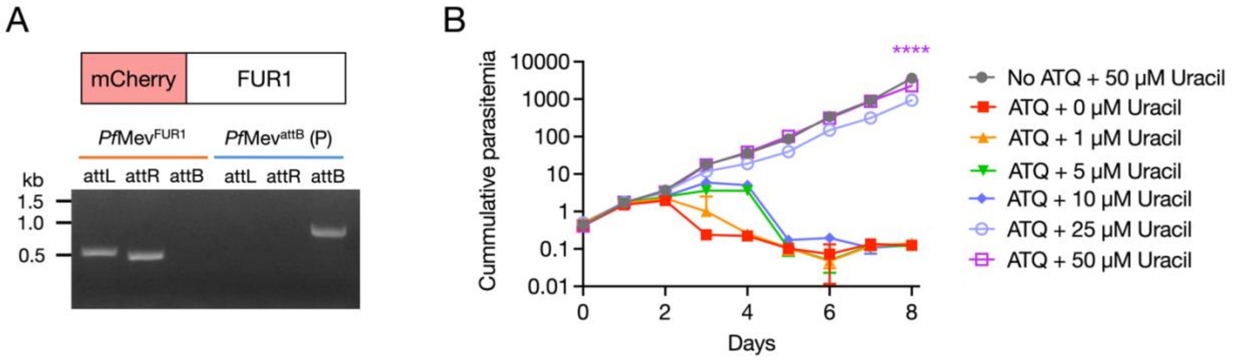


**Fig. S6. Parasites expressing yeast FUR1 are rescued from atovaquone treatment by uracil supplementation.** (A) Genomic integration of p15.mCherry-FUR1 in *Pf*Mev$^{attB}$ (Parent, 'P') parasites was confirmed by PCR amplification (*att*L and *att*R products). The intact P230p locus (*att*B product) was detected only in the parent (blue) and not in the transgenic line (orange). Refer to **Fig. S1A** for a detailed illustration of the integration of p15.mCherry plasmids into *att*B sites in the *P. falciparum* genome. Primer pairs for verifying integration of the p15.Cherry plasmid into the P230p locus are listed in **Fig. S1B** along with the expected sizes of the PCR products. (B) Parasites were treated with either 0 or 500 nM atovaquone (ATQ) in combination with increasing concentrations of uracil. Parasitemia was quantified daily by flow cytometry, with 1:10 dilutions performed every two days to prevent overgrowth. Error bars indicate standard deviations from means. Two-way ANOVA was computed using GraphPad Prism 10, followed by Bonferroni's correction; ****, P ≤ 0.0001. P-value only shown for multiple comparison analysis between samples from ATQ + 50 µM Uracil and No ATQ + 50 µM control conditions. The addition of 50 µM uracil largely restores growth to parasites treated with ATQ.

A

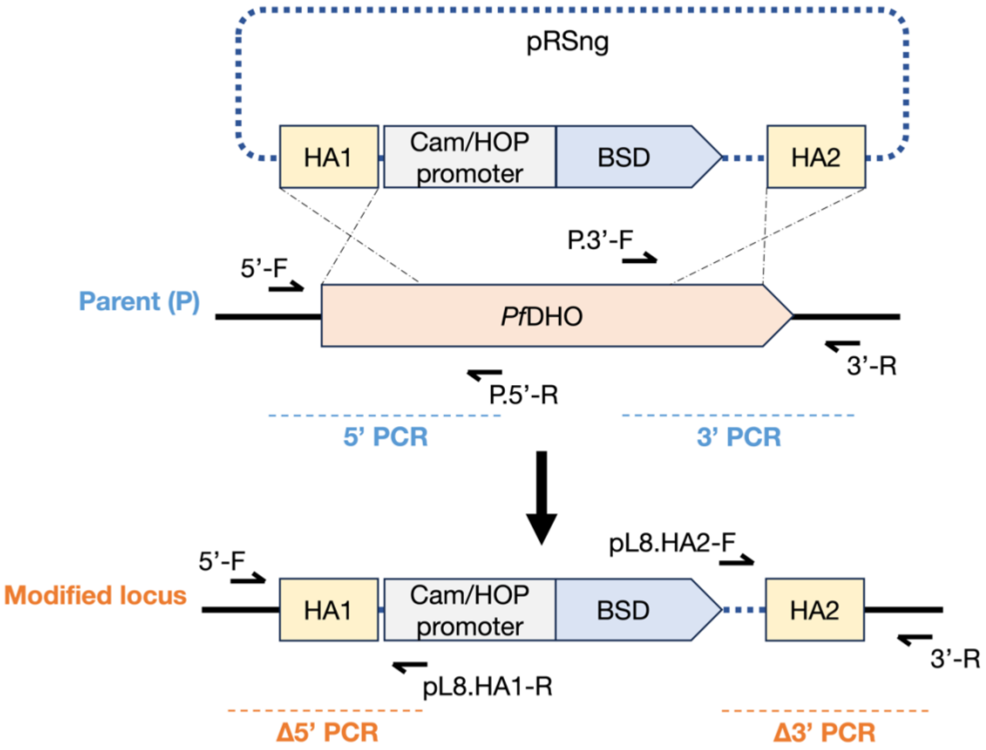


B

| Lane | PCR reaction | Primer Combination | | Product size (bp) | Parasite Line |
|---|---|---|---|---|---|
| 1 | Δ5' | DHO.5'-F<br>Pyr_057 | pL8.HA1-R<br>Pyr_062 | 619 | Δ*Pf*DHO |
| 2 | Δ3' | pL8.HA1-F<br>Pyr_061 | DHO.3'R<br>Pyr_058 | 391 | |
| 3 | 5' | DHO.5'-F<br>Pyr_057 | DHO.P.5'-R<br>Pyr_060 | – | |
| 4 | 3' | DHO.P.3'-F<br>Pyr_059 | DHO.3'-R<br>Pyr_058 | – | |
| 5 | Δ5' | DHO.5'-F<br>Pyr_057 | pL8.HA1-R<br>Pyr_062 | – | *Pf*Mev$^{FUR1}$ Parent (P) |
| 6 | Δ3' | pL8.HA1-F<br>Pyr_061 | DHO.3'R<br>Pyr_058 | – | |
| 7 | 5' | DHO.5'-F<br>Pyr_057 | DHO.P.5'-R<br>Pyr_060 | 575 | |
| 8 | 3' | DHO.P.3'-F<br>Pyr_059 | DHO.3'-R<br>Pyr_058 | 645 | |

C

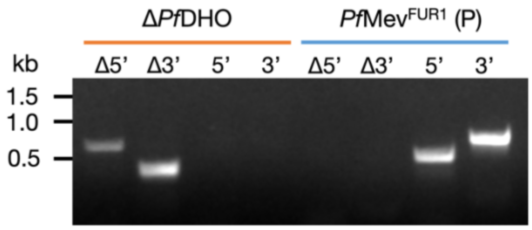

**Fig. S7. Schematic of CRISPR/Cas9-mediated disruption of *Pf*DHO.** (A) The pRSng plasmid encodes a blasticidin deaminase (BSD) expression cassette flanked by sequences with homology to *Pf*DHO, indicated by dotted lines. The Cam/HOP promoter drives expression of BSD. Arrows indicate the positions of primers used for diagnostic PCR. HA1 and HA2 are the homology arms utilized for recombination of pRSng into the *Pf*DHO locus in *Pf*Mev$^{FUR1}$ parasites (Parent, 'P'). (B) Primer pairs for verifying gene knockouts are listed along with the expected sizes of the PCR products. (C) The disruption of *Pf*DHO was confirmed by PCR amplification (Δ5' and Δ3' products). The intact *Pf*DHO locus (5' and 3' products) was detected only in the parent (blue) and not in the Δ*Pf*DHO line (orange).

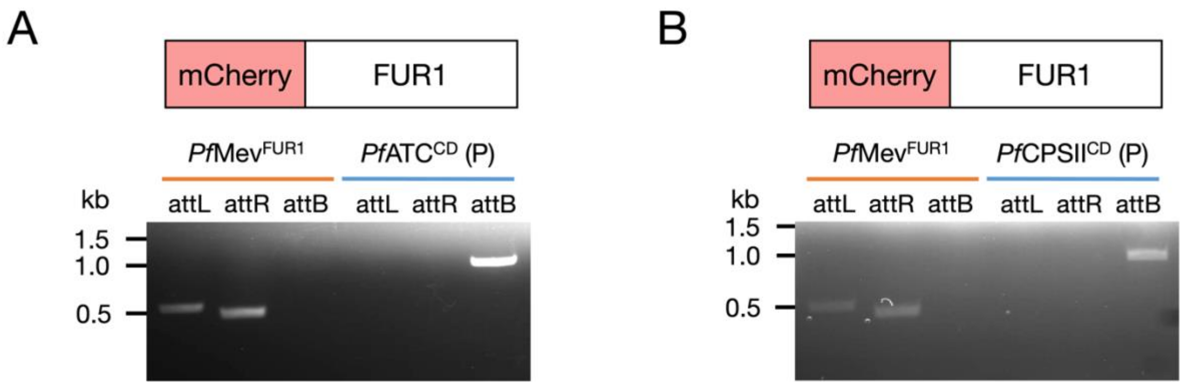


**Fig. S8. Bxb1-mediated integration of mCherry-FUR1 expression constructs into the *P. falciparum* genome.** Genomic integration of the p15.mCherry-FUR1 plasmid at the P230p locus in (A) *Pf*ATC$^{CD}$ (Parent, 'P') or (B) *Pf*CPSII$^{CD}$ (Parent, 'P') parasites was confirmed by PCR amplification of *att*L and *att*R products. The intact P230p locus (*att*B product) was detected only in the parent (blue) and not in the transgenic lines (orange). For a detailed schematic of p15.mCherry plasmid integration into *att*B sites, refer to **Fig. S1A**. Primer pairs for verifying integration into the P230p locus are listed in **Fig. S1B** along with the expected PCR product sizes.

A

| Lane | PCR reaction | Primer Combination x= CG6 | | Product size (bp) | Parasite Line |
|---|---|---|---|---|---|
| 1 | attL | x-F Pyr_009 | attB.int.5’ Pyr_011 | 334 | $NF54^{FUR1}$ |
| 2 | attR | attB.int.3’ Pyr_012 | x-R Pyr_010 | 1067 | |
| 3 | attB | x-F Pyr_009 | x-R Pyr_010 | – | |
| 4 | attL | x-F Pyr_009 | attB.int.5’ Pyr_011 | – | $NF54^{attB}$ Parent (P) |
| 5 | attR | attB.int.3’ Pyr_012 | x-R Pyr_010 | – | |
| 6 | attB | x-F Pyr_009 | x-R Pyr_010 | 1288 | |

B

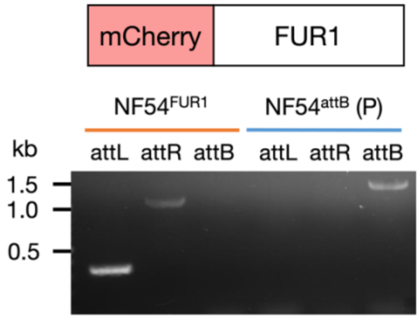


C

| Lane | PCR reaction | Primer Combination x= *Pf*DHODH | | Product size (bp) | Parasite Line |
|---|---|---|---|---|---|
| 1 | 5’ | x.5’-F Pyr_046 | Apt-5R Pyr_050 | 1001 | *Pf*DHODH$^{CD}$ |
| 2 | 3’ | Hsp86-F Pyr_049 | x.3’-R Pyr_047 | 507 | |
| 3 | P | x.5’-F Pyr_046 | x.3’-R Pyr_050 | – | |
| 4 | 5’ | x.5’-F Pyr_046 | Apt-5R Pyr_050 | – | $NF54^{FUR1}$ Parent (P) |
| 5 | 3’ | Hsp86-F Pyr_049 | x.3’-R Pyr_047 | – | |
| 6 | P | x.5’-F Pyr_046 | x.3’-R Pyr_050 | 1118 | |

D

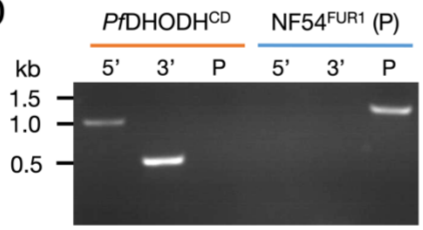

**Fig. S9. Generation of a *Pf*DHODH conditional knockdown line in NF54$^{attB}$ parasites expressing FUR1.** (A) Primer pairs for verifying p15.Cherry-FUR1 integration into the CG6 genomic locus of NF54$^{attB}$ parasites (Parent, ‘P’) are provided. (B) Genomic integration at the CG6 locus was confirmed by PCR (*att*L and *att*R products), with the intact CG6 locus (*att*B product) detected only in the parent line (blue) and not in the transgenic line (orange). Refer to **Fig. S1A** for a detailed schematic of p15.mCherry plasmid integration into *att*B sites. (C) Primer pairs for verifying integration of the pKD$^{tagless}$-*Pf*DHODH plasmid into the 3’ UTR of *Pf*DHODH are listed. (D) The genotype of the PDHODH$^{CD}$ parasite line was confirmed by PCR amplification of the 5’ and 3’ loci at the genomic insertion site (orange). The NF54$^{FUR1}$ parent (P) line served as a control (blue). Refer to **Fig. S3A** for a schematic of knockdown plasmid insertion into the 3’ UTR of target genes.

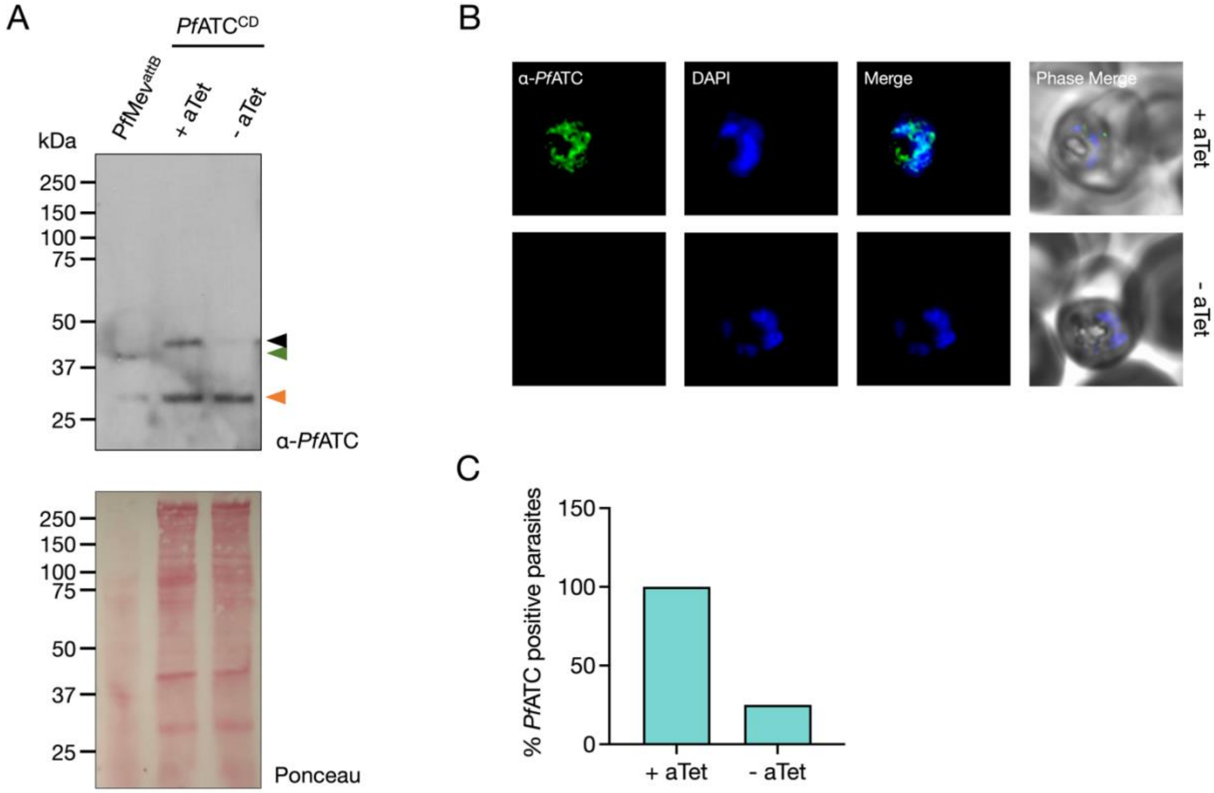


**Fig. S10. Validation of *Pf*ATC antibody specificity.** (A) Lysates from *Pf*Mev$^{attB}$ (control) and *Pf*ATC$^{CD}$ parasites were probed with α-*Pf*ATC. The antibody detected a ~40 kDa band in *Pf*Mev$^{attB}$ lysates (green arrow), consistent with the predicted molecular weight of *Pf*ATC (43.3 kDa). In *Pf*ATC$^{CD}$ parasites, a slightly higher band is observed (black arrow, +aTet condition) but absent when *Pf*ATC is depleted by removal of aTet from culture media (-aTet). This higher product is consistent with the predicted 45 kDa size of the endogenously C-terminally 2×FLAG tagged *Pf*ATC. The orange arrow marks a non-specific band. The blot was stained with Ponceau S to show relative loading. (B) *Pf*ATC$^{CD}$ parasites cultured under +aTet and -aTet conditions were fixed and probed with affinity-purified α-*Pf*ATC in an immunofluorescence assay. Representative images show specific staining by α-*Pf*ATC antibodies (green) in parasites grown under +aTet conditions, while no signal was detected in parasites grown under -aTet conditions. Images represent fields that are 10 µm long by 10 µm wide. (C) Fraction of *Pf*ATC$^{CD}$ parasites exhibiting α-*Pf*ATC staining under +aTet and -aTet conditions. All parasites in the +aTet condition were positive for α-*Pf*ATC staining (total cells imaged = 16), while the signal was markedly reduced in the -aTet condition (total cells imaged = 36).
