## Supplementary material for "Genetic analysis of pyrimidine biosynthetic enzymes in *Plasmodium falciparum*": File S2

### **Generation of homology arms (HA) for PfDHO.pTDN plasmid**

HA1: red
Recoded sequence: lowercase font
HA2: blue
▼ *Bsa*I cut site on + strand
▲ *Bsa*I cut site on - strand

#### HA2 + recoded HA1 Fragment synthesized by LifeSct LLC

GGCGCGCCTAAAACTCAAGCATTTTTACGATTATTTTTTTTTTATCTTTATTATTTTATTATTA
*Asc*I

TAATTAAATTATTATTATAATTAAATTATTATTATATTATAATTATATTATTATTATAATTATA

TTATTATTATTAGGATAATAATAATAATTATTATTATTTTTTTTTTAATATTTTTTAGTATGAT

TAATAATTATAAATATGTATATAATATATATATATACATACATATATATATTTTTTCATTATGT

GGCATTTCAAAAAAAAAATGAAGTATTTTATTTGTTTAATATATTTTTTATTTCATTAAATTAT

TATCATACGATATCCTCGAGGGATCCGAGACCAAGGCCTTGGTCTCGTCCC**aTTccTtGCtGGa**
*Eco*RV *Bam*HI *Bsa*I *Bsa*I **recoded gRNA**

**AAaACac**TtGATTATGAtATACATTATGTtAGTAAGTTTGACGTCTCCGGAGATTATAAAGACC
**sequence** *Aat*II *Bsp*EI

ATGATGGAGATTATAAAGATCACGATATAGATTACAAAGACCATGATAGTTAAGGGCCC
3xFLAG *Psp*OMI

#### HA1 PCR amplification from P. falciparum genomic DNA

Forward primer DHO.HA1-F:

GGTGGTCTCGGATCCGAAATGTAGCAGGTTCCATAACACC
*Bsa*I

Reverse primer DHO.HA1-R:

GGTGGTCTCGGGGACTACACCATTATATTCTCTTGG
*Bsa*I

HA1 PCR product:

GGTGGTCTCGGATCCGAAATGTAGCAGGTTCCATAACACCTCATCATTTATATTTAACAATAGA
*Bsa*I

TGATGTTGTTAATATGGATATATATGATCATGCAATAGATAACACATATATTGAAAAATATATA

AAAAATACATATCATTATTGTAAGCCATTACCAAAATTGTTAGAAGATAAAATTGCTTTACAAG

ATGTTATAAAAGATGATTTTCCAAGAGTCTTTTTGGGTTCGGATTCAGCACCTCATTACAAAGT

TATGAAGCGCAAACCCTACTATAAACCAGGAATATACACACAACCATTTTTAATAAATTATGTT

GCTCATATATTGAACAAATTCGATGCTTTAGATAAGATGGAAAATTTTACCTCAAAAAATGCTT

CCCTCTTTCTAAATTTAGCAGAAAAAAAAAAAATTGGCAAAATATTACATATGTGTGGAAAAACA

TCCATTTAAATTACCAAGAGAATATAATGGTGTAGTCCCCGAGACCACC

*Bsa*I

The synthesized fragment and the HA1 PCR product were digested with BsaI to generate complementary sticky ends, followed by ligation to generate the *Asc*I-HA2-*Eco*RV-*Bam*HI-HA1-3xFLAG-*Psp*OMI construct below.

GGCGCGCCTAAAACTCAAGCATTTTTACGATTATTTTTTTTTATCTTTATTATTTTATTATTA

*Asc*I

TAATTAAATTATTATTATAATTAAATTATTATTATATTATAATTATATTATTATTATAATTATA

TTATTATTATTAGGATAATAATAATAATTATTATTATTTTTTTTTTAATATTTTTTAGTATGAT

TAATAATTATAAATATGTATATAATATATATATATACATACATATATATATTTTTTCATTATGT

GGCATTTCAAAAAAAAAATGAAGTATTTTATTTGTTTAATATATTTTTTATTTCATTAAATTAT

TATCATACGATATCCTCGAGGGATCCCGAAATGTAGCAGGTTCCATAACACCTCATCATTTATA

*Eco*RV *Bam*HI

TTTAACAATAGATGATGTTGTTAATATGGATATATATGATCATGCAATAGATAACACATATATT

GAAAAATATATAAAAAATACATATCATTATTGTAAGCCATTACCAAAATTGTTAGAAGATAAAA

TTGCTTTACAAGATGTTATAAAAGATGATTTTCCAAGAGTCTTTTTGGGTTCGGATTCAGCACC

TCATTACAAAGTTATGAAGCGCAAACCCTACTATAAACCAGGAATATACACACAACCATTTTTA

ATAAATTATGTTGCTCATATATTGAACAAATTCGATGCTTTAGATAAGATGGAAAATTTTACCT

CAAAAAATGCTTCCCTCTTTCTAAATTTAGCAGAAAAAAAAAAAATTGGCAAAATATTACATATG

TGTGGAAAAACATCCATTTAAATTACCAAGAGAATATAATGGTGTAGTCCCC**aTTccTtGCtGG**

**recoded gRNA**

**aAAaACac**TtGATTATGAtATACATTATGTtAGTAAGTTTGACGTCTCCGGAGATTATAAAGAC

**sequence** *Aat*II *Bsp*EI

CATGATGGAGATTATAAAGATCACGATATAGATTACAAAGACCATGATAGTTAAGGGCCC

3xFLAG *Psp*OMI

This ligation product was then cloned into the *Asc*I and *Psp*OMI sites of the pTDN vector.

#### Synthesized TetR-DOZI sequence for pTDN plasmid

*Avr*II
<u>CCTAGG</u>ATGAGTAGATTAGATAAAAGTAAAGTGATTAACAGTGCATTAGAGTTACTTAATGAGG
TAGGAATAGAAGGTTTAACAACCCGTAAATTAGCCCAGAAGTTAGGTGTAGAGCAGCCTACATT
GTATTGGCATGTAAAAAATAAGAGAGCTTTGTTAGACGCCTTAGCCATTGAGATGTTAGATAGG
CACCATACTCACTTCTGCCCTTTAGAAGGTGAAAGTTGGCAAGATTTTTTACGTAATAACGCTA
AAAGTTTTAGATGTGCTTTATTAAGTCATAGAGATGGAGCAAAAGTACATTTAGGTACAAGACC
TACAGAAAAACAGTATGAAACTTTAGAAAATCAATTAGCCTTTTTATGCCAACAAGGTTTTTCA
TTAGAGAACGCATTATATGCTTTAAGTGCTGTGGGGCATTTTACCTTAGGTTGCGTATTGGAAG
ATCAAGAGCATCAAGTTGCTAAAGAAGAAAGGGAAACACCTACTACTGATAGTATGCCTCCATT
ATTACGACAAGCTATTGAATTATTTGATCACCAAGGTGCAGAGCCAGCCTTCTTATTCGGACTT
GAATTGATTATATGCGGATTAGAAAAACAACTTAAATGTGAAAGTGGGTCT<u>ACTAGT</u>AGTTATA
AAACAAATTGTACGAACTCTAATGCTAATACAAATACTTTGAATAGTTCTTCAAATTATAACAA
AATAGATGATAATATAATATTAGATGAAGAATGGAAAAAGAAAATTCTGGAACCATTAAAAGAT
TTAAGATATAAGACAGAAGATGTAACGAAAACGAAAGGCAATGAATTTGAAGATTATTTTTTGA
AGAGAGAATTATTAATGGGTATCTTTGAAAAAGGATATGAGAAACCATCACCTATACAAGAGGA
AAGTATACCTGTAGCTTTGGCTGGAAAAAATATTTTAGCAAGGGCAAAAAATGGTACCGGCAAA
ACAGCAGCTTTTGCTATACCCTTACTAGAGAAATGTAATACCCACAAAAATTTTATTCAAGGAC
TCATTTTAGTACCCACGCGAGAACTTGCCCTACAGACCTCTGCTATGATTAAGGAATTAGGAAA
ACACATGAAAGTACAGTGTATGGTAACAACTGGTGGTACATCATTAAGAGAAGATATAATGAGG
TTGTATAATGTAGTTCATATTTTATGTGGTACTCCAGGAAGAATATTAGACTTAGCAAATAAGG
ATGTAGCAAATTTATCAGGTTGTCATATTATGGTTATGGATGAAGCAGATAAATTATTATCACC
TGAATTTCAACCTATAGTAGAAGAACTAATGAAATTTTTACCAAAAGAAAAGCAGATACTTATG
TATTCTGCTACCTTTCCTGTGACTGTAAAAGAATTTCGAGCTATTTATTTATCAGATGCCCATG
AAATAAATCTTATGGATGAATTAACCTTAAAAGGAATAACACAATATTATGCTTTTGTTAAAGA
AAGACAAAAAGTACATTGTTTAAATACATTATTTGCTAAACTTCAAATTAATCAAGCTATCATC
TTCTGTAATAGTATTACTAGGGTAGAACTACTAGCCAAAAAAATTACCGAACTAGGATATAGCT
CTTTTTACATTCATGCAAGAATGTCACAAACACATCGTAATCGTGTTTTCCATGATTTTAGAAA
TGGAGCATGTAGATGTTTAGTTTCATCAGATTTATTCACAAGAGGTATCGACATACAGTCAGTC
AATGTTGTTATCAATTTTGATTTCCCAAAAAATTCTGAAACTTATTTACATAGAATAGGAAGAT
CAGGAAGATACGGACATCTAGGACTAGCTATTAATCTTATAACTTTTGAAGATCGTTTTAATTT
ATATAAAATAGAAGTAGAACTAGGAACGGAAATACAACCAATACCAAACGAAATTGACCCATCC
TTATATACC<u>GCTAGC</u>
*Nhe*I

#### **Synthesized Neomycin sequence for pTDN plasmid**

*Ngo*MIV

GCCGGCATGAGCGCTATTGAACAAGATGGATTGCACGCAGGTTCTCCTGCTGCTTGGGTGGAAA
GACTATTTGGTTATGATTGGGCACAACAGACAATAGGATGCAGTGATGCAGCAGTATTTAGATT
ATCAGCTCAAGGAAGGCCGGTTCTTTTTGTAAAAACAGACTTATCCGGTGCATTAAATGAATTG
CAAGACGAAGCAGCACGATTATCGTGGTTAGCTACGACAGGTGTACCTTGTGCAGCTGTATTAG
ATGTTGTAACTGAAGCAGGAAGGGATTGGCTGCTATTGGGAGAAGTTCCTGGACAAGATTTATT
ATCATCTCATTTAGCTCCAGCCGAAAAAGTTAGTATAATGGCTGATGCAATGAGGAGATTACAT
ACTTTAGATCCAGCTACATGTCCATTTGATCATCAAGCTAAACATCGTATTGAGCGAGCACGTA
CAAGAATGGAAGCAGGTTTAGTTGATCAAGATGATTTAGATGAAGAACATCAAGGTTTAGCACC
AGCCGAATTATTTGCGAGGCTTAAAGCGAGAATGCCAGATGGTGATGATTTAGTCGTAACTCAT
GGGGATGCCTGTTTGCCTAATATAATGGTTGAAAATGGTAGATTTAGTGGATTTATTGATTGTG
GCAGACTAGGAGTGGCTGATAGATACCAAGACATAGCTTTAGCTACCAGAGATATTGCTGAAGA
ATTAGGTGGGGAATGGGCTGATCGCTTCCTCGTACTTTATGGAATCGCCGCACCCGATTCACAA
AGAATAGCTTTTTATAGATTATTAGATGAATTTTTCTAACCGGT

*Age*I
